# Transient SUMOylation Inhibition In Human Pre-adipocytes Stably Imprints a Transcriptional Beiging Fate

**DOI:** 10.1101/2025.03.07.642034

**Authors:** Patrizia Maria Christiane Nothnagel, Paul-Arthur Meslin, Jonas Aakre Wik, Yunna Erika Strøm, Magnar Bjørås, Jorrit Martjin Enserink, Bjørn Steen Skålhegg, Nolwenn Briand, Anthony Mathelier, Pierre Chymkowitch

**Author notes:** Present address: Department of Molecular Biology, Faculty of Science, Radboud Institute for Molecular Life Sciences, Oncode Institute, Radboud University Nijmegen, 6525 GA Nijmegen, The Netherlands.

## Abstract

SUMOylation regulates chromatin states and transcriptional programs that preserve cellular identity, yet how transient perturbation of the SUMO pathway impacts adipocyte plasticity remains unclear. Here we show that brief pharmacologic inhibition of SUMO conjugation in human pre-adipocytes using TAK-981 primes stable de novo beige differentiation in the presence of the PPARG agonist rosiglitazone. Transient TAK-981 exposure before adipogenic induction produces long-lasting changes in the transcriptome and metabolism of mature adipocytes, including robust induction of canonical beiging markers like *UCP1* and increased mitochondrial respiration. Mechanistically, ATAC-seq and transcription factor footprinting revealed immediate and durable chromatin remodeling and early mobilization of CEBP family members, followed by stable activation of CEBPA and PPARG regulatory networks. ChIP experiments demonstrated loss of H3K27me3 and gain of H3K27ac at PPAR response elements within key thermogenic enhancers, with increased PPARG occupancy across the *UCP1* regulatory unit. This mechanism is enforced by enhanced cAMP-PKA-p38 signaling, and stabilization of beiging transcriptional activators. Our data support a model in which transient relief of SUMO-mediated repression unlocks dominant regulatory units, notably the *UCP1* enhancer cluster, producing a monomorphic, cell type specific reprogramming toward adaptive thermogenesis. These findings identify SUMOylation as a reversible epigenetic barrier to adipocyte beiging and suggest that temporally controlled SUMO pathway inhibition combined with PPARG activation could be exploited to modulate adipose tissue thermogenic capacity.

**Graphical abstract:** 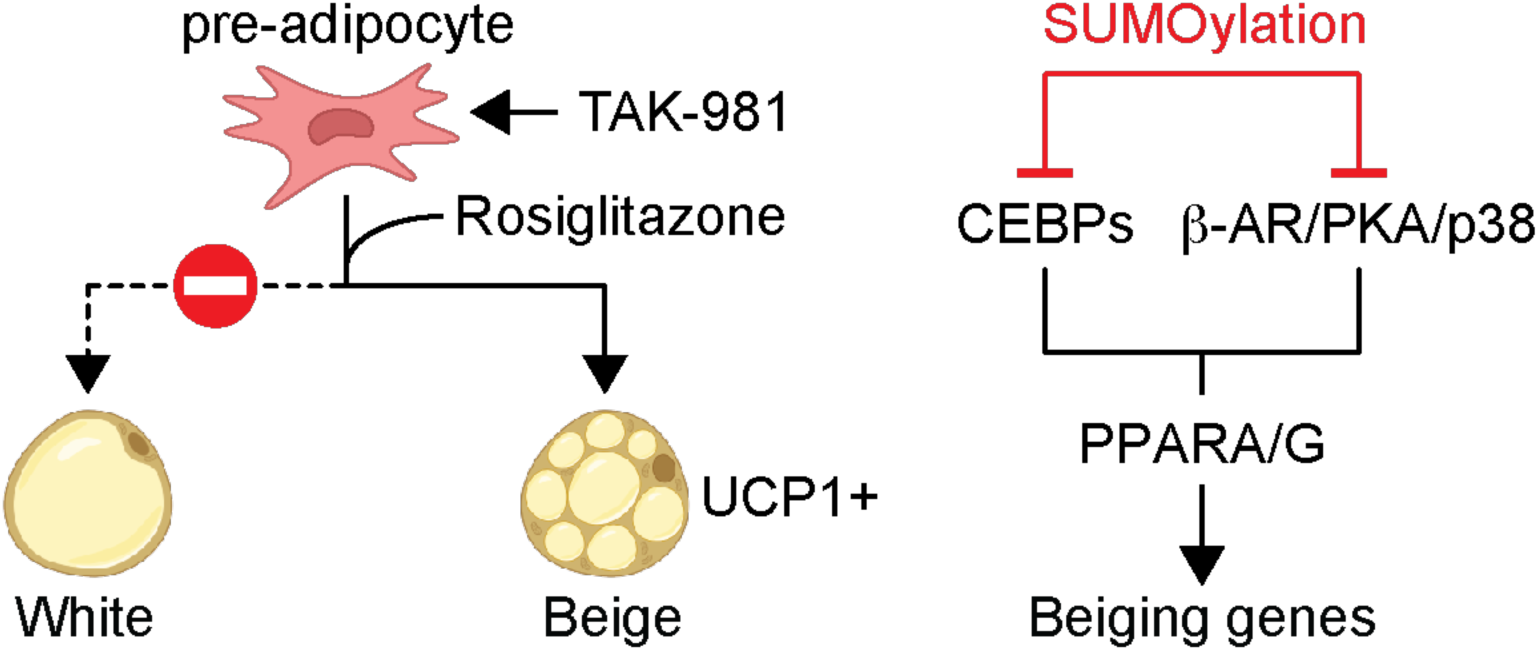

SUMOylation restricts *de novo* differentiation of pre-adipocytes into beige adipocytes by repressing cAMP signaling and preventing epigenetic and transcriptional reprogramming of white adipocytes.

## Introduction

Post-translational modification by the small ubiquitin-like modifier (SUMO) is a reversible process in which a SUMO peptide is covalently conjugated to target proteins, thereby modulating their function, localization, physical interactions and stability (1). Over the past decade, extensive research investigated the role of the SUMOylation pathway in cellular differentiation and development, employing diverse *in vitro* and *in vivo* models (2–5). Although the precise molecular mechanisms underlying SUMOylation and its substrate specificity remain incompletely defined, accumulating evidence underscores its indispensability in developmental processes and its involvement in nearly all cellular functions, particularly nuclear processes. Notably, a dynamic equilibrium between SUMOylation and deSUMOylation is fundamental for the formation and functional maturation of multiple cell types and organs, including adipose tissue (4–13).

Crucially, SUMOylation has been implicated in the repression of cellular reprogramming through modification of key regulators of chromatin architecture, epigenetic landscapes, and transcriptional activity (2,3,5,14,15). These findings highlight the essential role of SUMOylation in maintaining epigenetic integrity and cellular homeostasis. Unsurprisingly, dysregulation of this pathway has been linked to pathological conditions, including oncogenesis (16).

Adipogenesis is an excellent model system for studying the role of SUMOylation in epigenetic memory and cellular reprogramming. Notably, the formation and functional maturation of white adipose tissue (WAT) appear to be dependent on a dynamic SUMOylation/deSUMOylation cycle, which orchestrates gene expression and repression programs essential for adipogenic differentiation (5,10,17). WAT plays a central role in energy homeostasis by regulating lipid storage and mobilization in response to metabolic demands, which has been linked to SUMOylation (18). Under conditions of positive energy balance, WAT accumulates excess lipids to prevent lipotoxicity and associated metabolic complications, such as hepatic steatosis. Conversely, during energy deficit, WAT undergoes lipolysis, releasing fatty acids as an alternative energy source to maintain systemic energy balance.

Remarkably, WAT exhibits plasticity and undergoes phenotypic conversions in response to environmental stimuli such as cold exposure. This process, termed adaptive thermogenesis or “beiging” is characterized by the emergence of beige adipocytes within WAT (19,20). Unlike classical white adipocytes, which primarily function as lipid storage depots, beige adipocytes possess the capacity to oxidize fatty acids and utilize oxygen to generate heat through uncoupling of the mitochondrial respiratory chain, a process mediated by uncoupling protein 1 (UCP1) (21).

Although white and beige adipocytes exhibit largely overlapping gene expression profiles (22–25), beige adipocytes uniquely express genes associated with mitochondrial uncoupling and energy dissipation, including *UCP1* (19). Evidence suggests that beiging occurs through the transdifferentiation of mature white adipocytes into beige adipocytes in response to hypothermic conditions, a process that is reversible in a temperature dependent fashion (26). Transcriptomic analyses further support this model, revealing that a subset of white adipocytes may be transcriptionally primed for beige conversion (20). In addition to transdifferentiation, beige adipocytes may arise from distinct progenitor populations or via the reprogramming of differentiating white adipocytes (27,28). However, the molecular mechanisms governing the induction or suppression of white-to-beige gene expression programs and associated epigenetic remodeling remain largely unresolved. Elucidating these mechanisms may provide critical insight into the regulation of adaptive thermogenesis and offer novel therapeutic strategies for combating obesity (29).

Globally, obesity and its related conditions, including type 2 diabetes, now cause more deaths than starvation (The Lancet, Global Burden of Disease)^1^. Strong evidence links obesity to a higher risk of several cancers (30–33) and severe SARS-CoV-2 infection, with severity correlating with body mass index (34). Current treatments like diets, medications, exercise, surgery, and liposuction often fail to restore metabolic balance or sustain weight loss, possibly due to an epigenetic obesogenic memory (21,35). Understanding WAT “beiging” and its role in energy expenditure could offer promising strategies to combat obesity and its complications (19,29,36).

Given its established role in maintaining cellular identity, we hypothesized that SUMOylation may play a regulatory role in limiting the plasticity of white adipose tissue (WAT), specifically the transcriptional and metabolic reprogramming associated with the conversion of white adipocytes to beige adipocytes. In this study, we demonstrate that transient inhibition of SUMOylation using TAK-981 in human pre-adipocytes is sufficient to induce a stable chromatin, epigenetic and transcriptional white-to-beige adipocyte transition. This conversion involved activation of the thermogenic pathway and is accompanied by mitochondrial uncoupling and increased energy expenditure, a process that is dependent on the peroxisome proliferator-activated receptor gamma (PPARG) agonist rosiglitazone.

## Materials and Methods

### Cell culture, differentiation and treatments

Human white pre-adipocytes (hTERT A41hWAT-SVF, ATCC, CRL-3386) were maintained and differentiated based on previously established protocols (37). Cells were maintained in StableCell^TM^ DMEM High Glucose Medium (Sigma, D0819), supplied with 10% Fetal Bovine Serum (FBS, F7524) and 1% Penicillin/Streptomycin (PenStrep, Gibco, 15140-122) at 37°C in a humidified incubator at 5% CO_2_ in air atmosphere.

Pre-adipocytes were grown to full confluence and kept at full confluence for 2 days. Next, differentiation was induced by adding StableCell^TM^ DMEM High Glucose Medium, supplied with 2% FBS, 1% PenStrep, 1 µM Dexamethasone (D4902, Sigma), 0.5 mM 3-Isobutyl-1-methylxanthine (I5879, Sigma), 0.1 µM human insulin (91077C, Sigma), 17 µM Pantothenate (P5155, Sigma), 22 µM Biotin (B4639, Sigma) and 1 µM Rosiglitazone (R2408, Sigma) for 7 days. On Day 7 (D7), the induction medium was replaced with the maintenance medium, consisting of StableCell^TM^ DMEM High Glucose Medium, supplied with 2% FBS, 1% PenStrep, 1 µM Dexamethasone, 0.1 µM human insulin, 17 µM Pantothenate and 22 µM Biotin for at least 14 days. The medium was refreshed every second or third day. The SUMOylation inhibitor TAK-981 (0.5 µM; Takara) was added to the cells at the indicated days for 48hrs (Fig. 1A). After the treatment, the cells were washed with phosphate-buffered saline (PBS) and differentiation was initiated as indicated (Fig. 1A). The PKA inhibitor Rp-8-CPT-cAMPS (500µM; C011, Biolog) was added for 5 minutes on D24 of differentiation. The cells were then washed twice in cold PBS and immediately used for sample preparation as described in Protein extraction and western blotting.

**Fig. 1:**
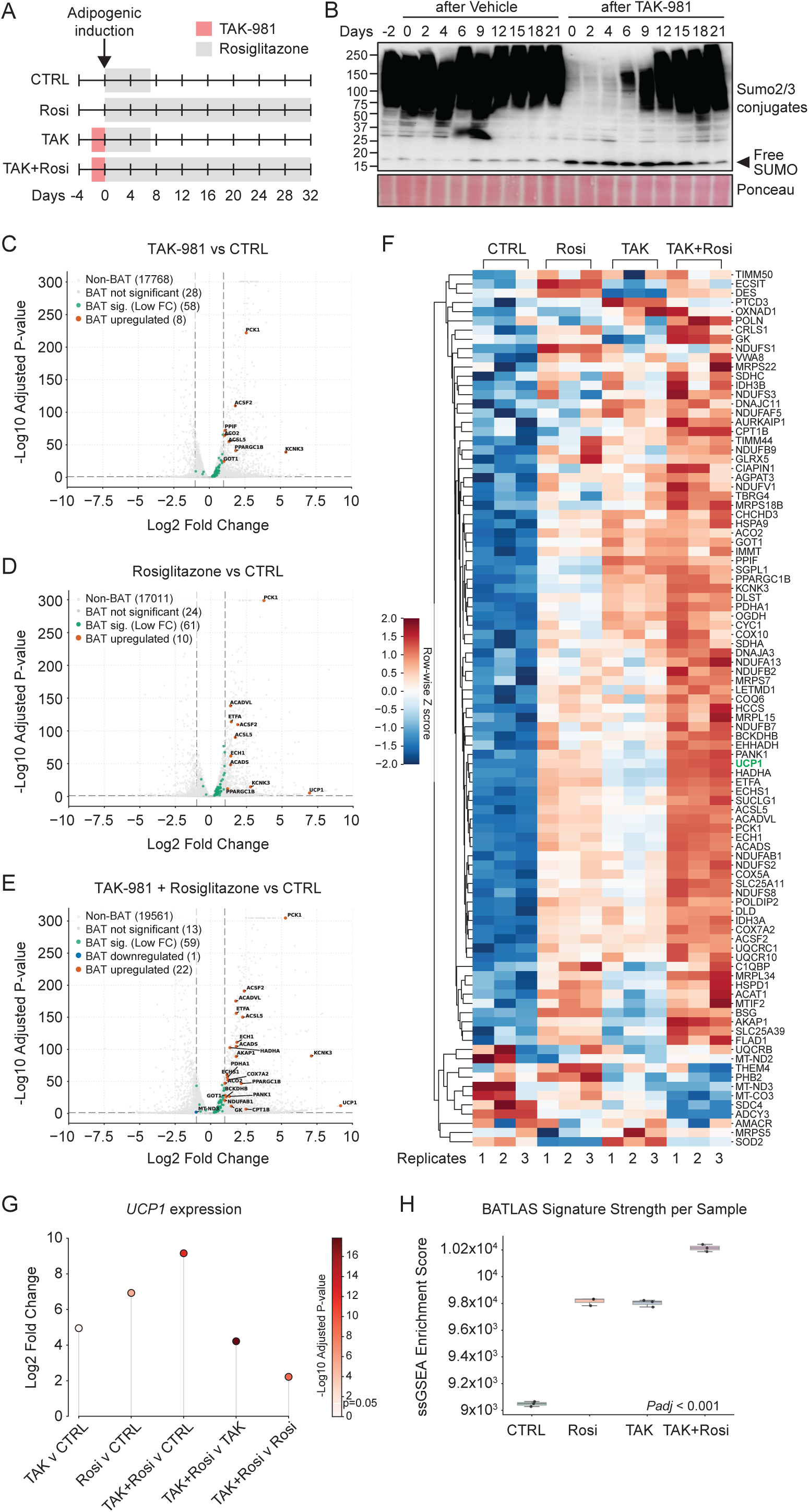
Analysis of the impact of transient SUMOylation inhibition in pre-adipocytes and rosiglitazone on gene expression in mature adipocytes. **(A)** Schematic representation of the experimental setup. Confluent human white pre-adipocytes were treated transiently for 48 hours with TAK-981 prior to adipogenic induction on day 0. Differentiation took place as described in the Methods with Rosiglitazone administration for the initial 7 days (CTRL and TAK) or for the complete period of differentiation (Rosi and TAK+Rosi). **(B)** Validation of transient SUMOylation inhibition by SUMO2/3 western blot at the indicated time points. **(C-E)** Volcano plots displaying differentially expressed genes (DEGs; log₂FC > 1 or log₂FC <-1; padj < 0.05) resulting from TAK-981 treatment of pre-adipocytes *(C)* rosiglitazone treatment *(D)* and co-treatment with TAK-981 + Rosiglitazone *(E)* compared to control cells. Data were collected in mature adipocytes 21 days post adipogenic induction. Significant BATLAS DEGs are highlighted. **(F)** Heatmap visualization of the expression patterns of BATLAS genes across all conditions described in Fig 1A. Biological replicates were ordered by experimental condition (CTRL, Rosi, TAK, TAK + Rosi) to facilitate comparisons. **(G)** Dot plot showing the expression of *UCP1* (RNA-seq) across conditions. **(H)** Enrichment of the BATLAS signature withing individual samples and condition assessed using ssGSEA. Statistical differences between experimental groups were evaluated using a one-way Analysis of Variance.

Human brown pre-adipocytes (hTERT-A41hBAT-SVF, ATCC, CRL-3385) were differentiated based on previously established protocols (38). SVF cells were seeded and grown to full confluency as described above. After two days at full confluency, the induction medium was added (StableCell^TM^ DMEM High Glucose Medium, supplied with 2% FBS, 1% PenStrep, 2µg/ml Dexamethasone, 0.5 mM 3-Isobutyl-1-methylxanthine, 5 µg/ml human insulin, 1 µg/ml Rosiglitazone and 50 nM T3. Differentiation was induced for two days, followed by changing to differentiation medium (StableCell^TM^ DMEM High Glucose Medium with 3-Isobutyl-1-methylxanthine, human insulin, Rosiglitazone and T3 as before).

Primary human ASCs were isolated from subcutaneous fat obtained by liposuction from a female donor (age 45; BMI 20,9 kg/m^2^) (39–41), obtained after donor’s informed consent as approved by the Regional Committee for Research Ethics for Southern Norway with number REK 2013/2102. Liposuccion material was washed repeatedly in HBSS (Gibco) supplemented with 1% penicillin-streptomycin (Gibco) and 2.5 µg / ml fungizone (Gibco) then digested in DMEM/F12 containing 1% type II collagenase (Worthington) at 37°C for 50 min. Following centrifugation, the stromal vascular fraction was separated from mature adipocytes, washed, filtered through 100 µm strainers, and plated in DMEM/F12 (17.5 mM glucose) supplemented with 10% FBS, 1% penicillin/streptomycin, and 20 ng/ml FGF2 (proliferation medium). After 24 h, non-adherent cells were removed, and adherent ASCs were expanded under standard culture conditions. For differentiations following TAK administration (Fig. 2C, D, F), the same procedure was applied as described above. After differentiation induction for 7 days, the cells were kept in differentiation maintenance medium. For differentiation following forskolin treatment (Supp. Fig. 4D), the cells were expanded and differentiated as described before (42). On day 15, forskolin (50µM) was administered for 6hrs. After the treatment, the cells were washed in PBS and maintained in the differentiation maintenance medium for 18hrs and 66hrs.

**Fig. 2:**
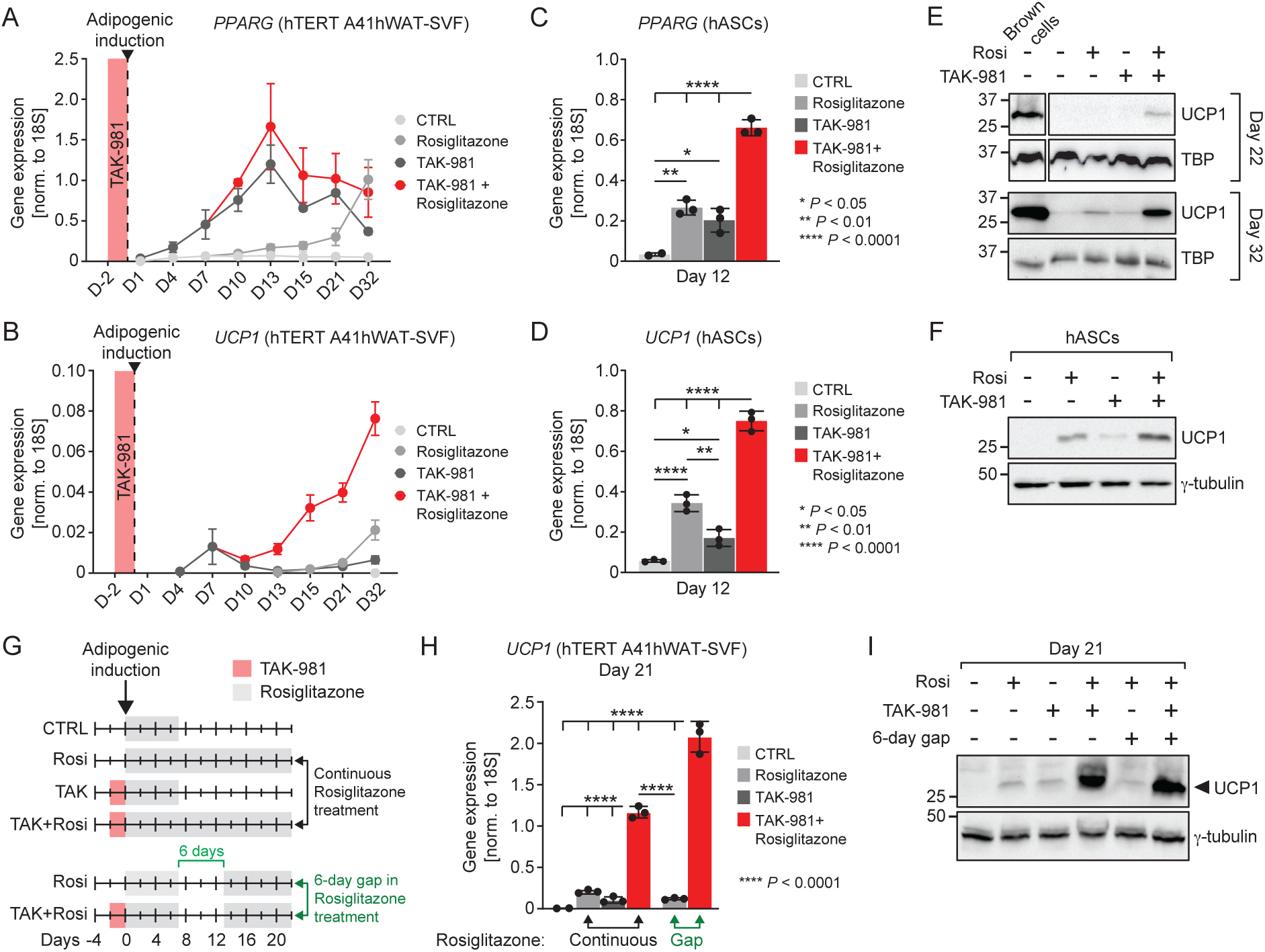
Analysis of TAK-981 and rosiglitazone effects on UCP1 expression in pre-adipocytes and human adipose stem cells. **(A)** Time-course analysis of *PPARG* mRNA expression in hTERT A4hWAT-SVF cells under the conditions described in Fig. 1A. Data were normalized to the expression of *18S* and represent the mean ± standard deviation from at least three independent experiments. **(B)** Time-course analysis of *UCP1* mRNA expression in hTERT A4hWAT-SVF cells under the same conditions as in *(A)*. Data were normalized to the expression of *18S* and represent the mean ± standard deviation from at least three independent experiments. **(C)** Analysis of *PPARG* mRNA expression in hASCs at day 12 post adipogenic induction under the same conditions as in *(A)*. Data were normalized to the expression of *18S* and represent the mean ± standard deviation from at least three independent experiments. **(D)** Analysis of *UCP1* mRNA expression in hASCs at day 12 post adipogenic induction under the same conditions as in *(A)*. Data were normalized to the expression of *18S* and represent the mean ± standard deviation from at least three independent experiments. **(E)** Western blot analysis of UCP1 protein levels at indicated days after adipogenic induction. Differentiated brown cells (hTERT A4hBAT-SVF) lysates were used as a positive control. **(F)** Western blot analysis of UCP1 protein levels in hASCs in the same conditions as in *(D)*. **(G)** Experimental conditions used in panels *(H)* and *(I)*. The upper section shows the conditions of continuous rosiglitazone treatment, as previously detailed in Fig. 1A. The lower section indicates the introduction of a six-day gap between differentiation induction with rosiglitazone and the prolongation of the treatment. **(H)** RT-qPCR analysis of *UCP1* mRNA expression in hTERT A4hWAT-SVF cells under the conditions in *(G)*. Data were normalized to the expression of *18S* and represent the mean ± standard deviation from at least three independent experiments. **(I)** Western blot analysis of UCP1 protein levels in conditions indicated in *(G)*. Statistical analyses: ONE-WAY-ANOVA with Tukey’s multiple comparison test.

**Fig. 3:**
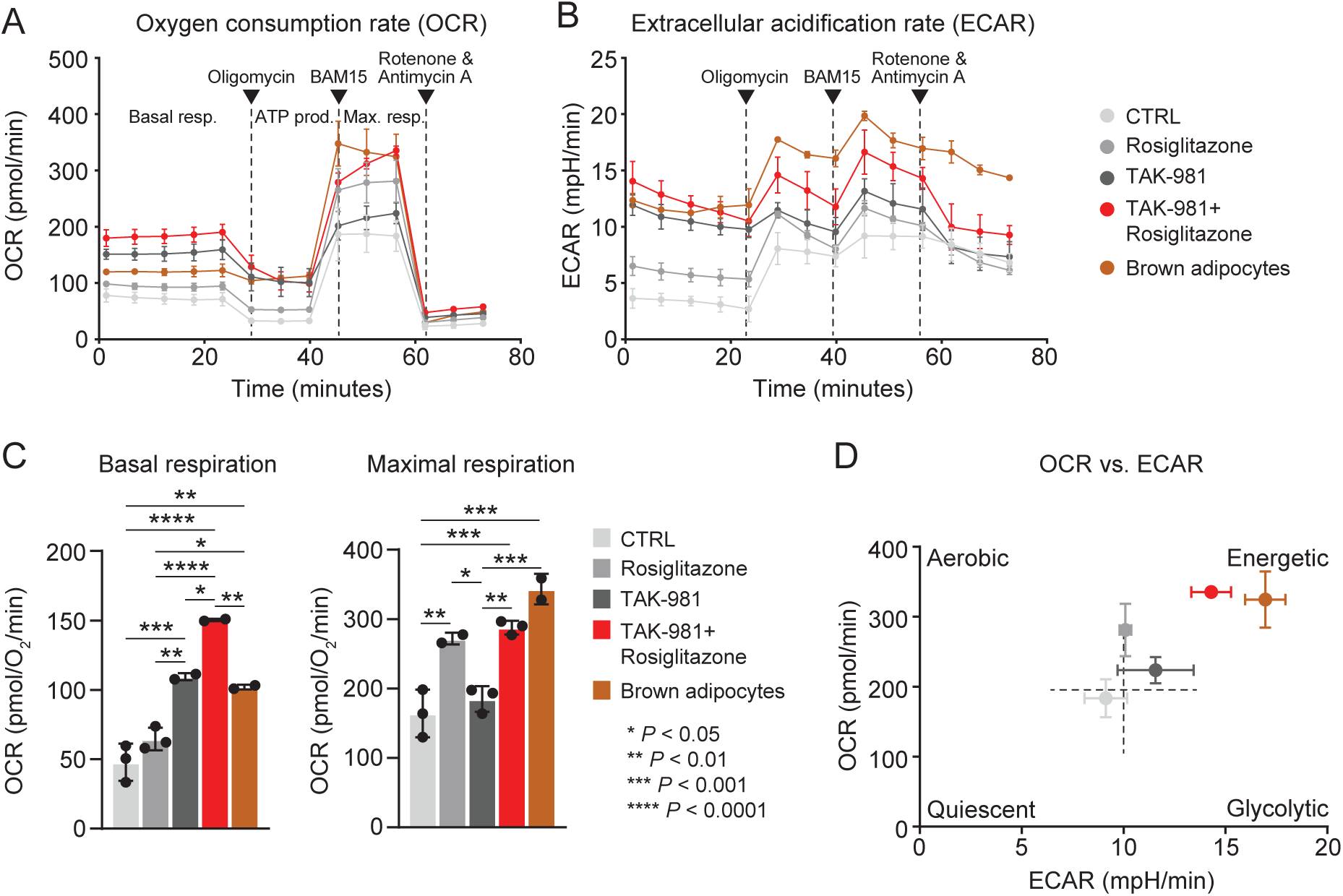
Metabolic assessment of TAK-981 and rosiglitazone effects on mitochondrial uncoupling. (A-B) Measurement of oxygen consumption rate (OCR) *(A)* and extracellular acidification rate (ECAR) *(B)* in mature adipocytes 22 days post-adipogenic induction. Differentiated brown cells (hTERT A4hBAT-SVF) were used as a positive control. **(C)** Basal and maximal respiration in mature adipocytes, 22 days post-adipogenic induction, were calculated after normalization to Crystal Violet staining. **(D)** Integration of OCR *(A)* and ECAR *(B)* data. Statistical analysis: one-Way-ANOVA followed by Tukey’s multiple-comparison test.

**Fig. 4:**
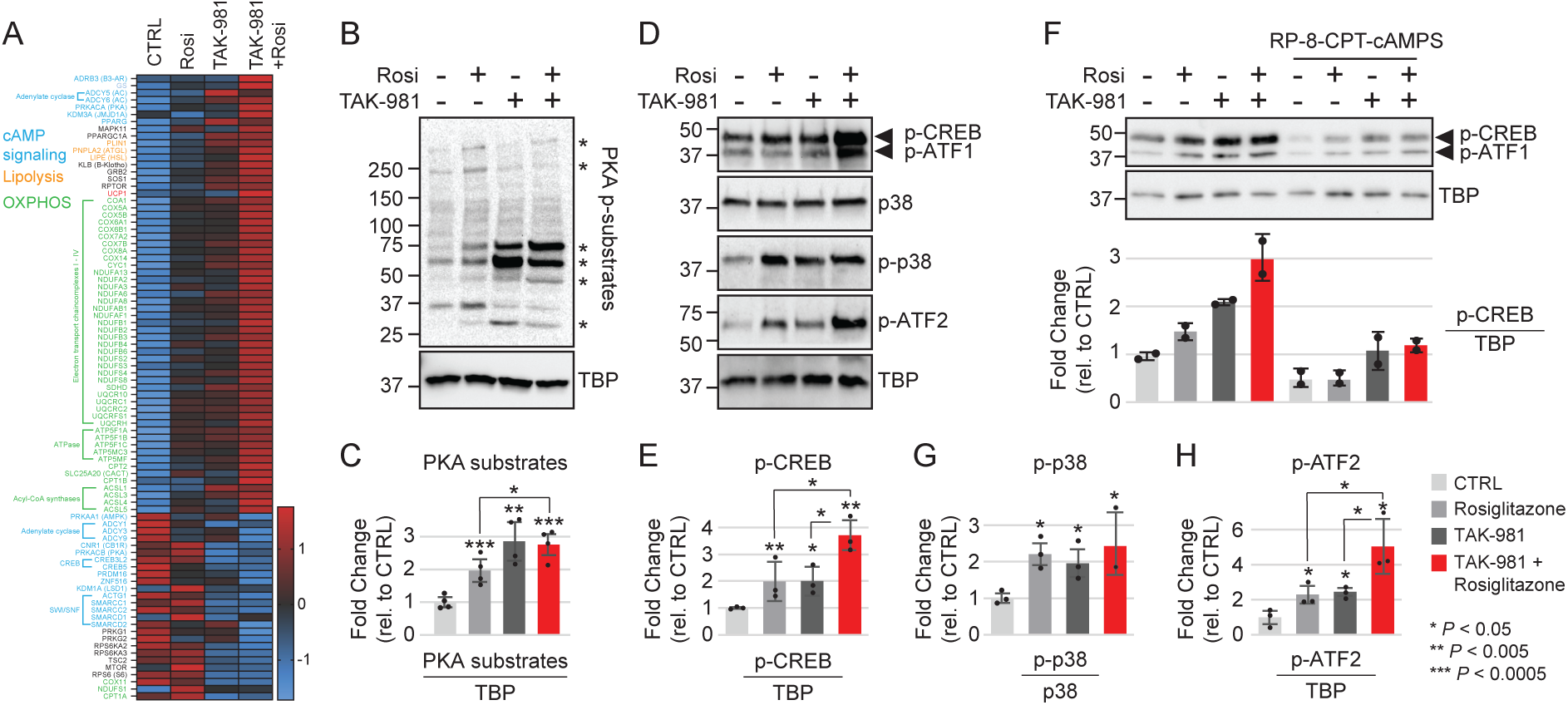
Effect of transient SUMOylation inhibition in pre-adipocytes on cAMP-PKA-p38 signaling and adaptive thermogenesis in mature adipocytes. **(A)** Heatmap showing the stable upregulation of most genes involved in adaptive thermogenesis and cAMP-PKA-p38 signaling 22 days after adipogenic induction. Z-scores were calculated and plotted in GraphPad Prism. Treatments of pre-adipocytes were performed as shown in Fig. 1A. **(B)** Western blot analysis of PKA substrates phosphorylation, assessed using a pan-PKA-target antibody, 22 days after adipogenic induction. TBP was used as a loading control. The asterisk (*) indicates substrates significantly affected by TAK-981 and/or rosiglitazone treatment. **(C)** Quantification of PKA substrate signals from *(B)*. The signal for PKA substrates and TBP was quantified using Fiji software, with PKA substrate signals normalized to TBP. Error bars represent the standard deviation of four independent experiments. Student’s T-test was used to calculate *P*-values. **(D)** Western blot analysis of p-CREB, p-ATF1, p38, p-p38, and p-ATF2, 22 days after adipogenic induction. TBP was used as a loading control. **(E)** Quantification of p-CREB / ATF-1 signals from *(D)*. Signals were normalized to TBP. Error bars represent the standard deviation of three independent experiments. Student’s T-test was used to calculate P-values. **(F)** Western blot analysis of p-CREB in absence or presence of the PKA inhibitor RP-8-CPT-cAMPS. TBP was used as a loading control. Quantification of p-CREB, normalized to TBP, is showed in the lower panel. Error bars represent the standard deviation of two independent experiments. **(G-H)** Quantification of p-p38 *(H)* and p-ATF2 *(I)* from experiment in *(F).* Signals were normalized to TBP for p-ATF2 and to p38 for p-p38. Error bars represent the standard deviation of three independent experiments. Student’s T-test was used to calculate *P*-values.

**Fig. 5:**
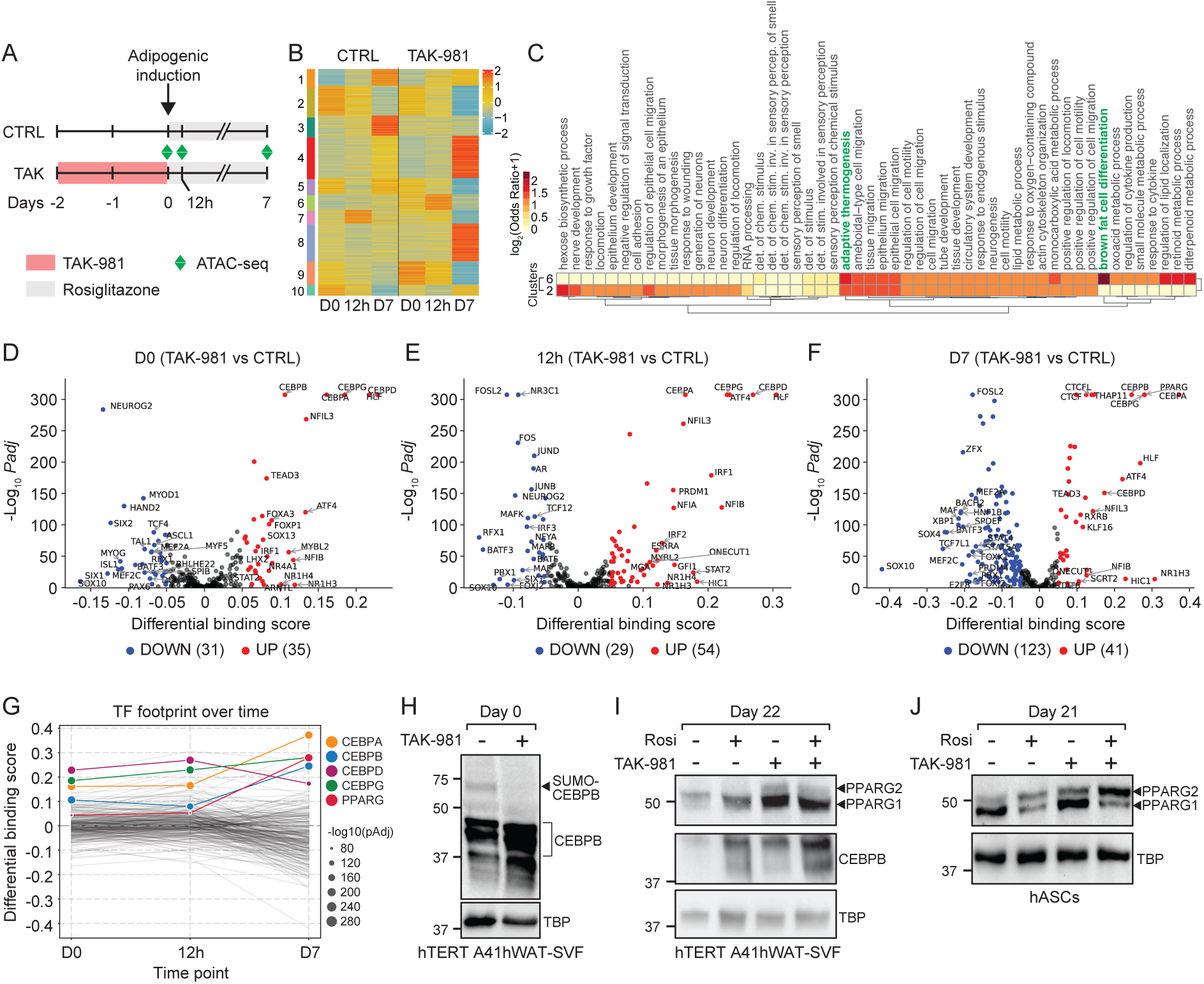
Stable mobilization of CEBPs and PPARG upon transient TAK-981 treatment in pre-adipocytes. **(A)** ATAC-seq experimental layout in hTERT A41hWAT-SVF pre-adipocytes. **(B)** Time series analysis and hierarchical clustering of significant variations in chromatin accessibility at day 0 (D0), 12 hours (12h) and day (D7) after TAK-981 treatment. **(C)** GOBP analysis of ATAC-seq clusters 2 and 6 revealed in *(B)*. See Suppl. Fig. 5A,B for an analysis of all clusters. **(D-F)** Volcano plots displaying the results of ATAC-seq TF footprint analyses performed at D0, 12h and D7 after TAK-981 or DMSO treatment. Colored dots indicate significant differentially mobilized TFs (TAK-981 vs DMSO). Differential binding score > 0.05 and *pAdj* < 0.001. **(G)** Time series analysis TF binding dynamics (TAK-981 vs DMSO) at D0, 12h and D7 after TAK-981 treatment. **(H)** Western blot analysis of CEBPB SUMOylation in DMSO and TAK-981-treated cells. **(I)** Western blot analysis of CEBPB and PPARG in DMSO, rosiglitazone, TAK-981-treated cells and co-treated hTERT A41hWAT-SVF pre-adipocytes. **(J)** Western blot analysis of PPARG in DMSO, rosiglitazone, TAK-981-treated cells and co-treated hASCs.

### Lipid staining

Differentiated adipocytes were washed with PBS and fixed in 10% formaldehyde for 45 minutes at room temperature. Cells were permeabilized with 60% isopropanol for 5 minutes at room temperature and stained with Oil Red O (Sigma-Aldrich, MAK194) for 15 minutes at room temperature. After 3 – 5 washes in mqH_2_O the staining was assessed under a Brightfield Microscope and a regular camera.

### Total RNA extraction

Cells were collected in TRIzol Reagent (Life Technologies, MAN0001271) and 200µL of chloroform was added per 1ml of TRIzol Reagent used. Samples were vortexed thoroughly and incubated on ice for 5 minutes before centrifugation at 13,200 RPM for 10 minutes at 4°C. The aqueous phase containing the RNA was collected and the RNA precipitated with 100% isopropanol and incubated on ice for 5 minutes. The RNA pellets were centrifuged at 13,200 RPM for 10 minutes at 4°C. The supernatant was removed and the RNA pellet washed in 70% ethanol, followed by incubation on ice for 5 minutes. After centrifugating the RNA at 13,000 RPM for 5 minutes at 4°C the supernatant was removed, and the RNA pellet briefly air dried.

Finally, the RNA was resuspended in RNAse-free H_2_O and incubated on ice for 20 minutes. RNA concentrations were determined using Nanodrop 3000.

### Reverse transcription and RT-qPCR

Total RNA was reverse transcribed using the QuantiTect Reverse Transcription Kit according to manufacturer’s instructions (Qiagen, 205313). RT-qPCR was performed using the HOT FIREPol® EvaGreen® qPCR Supermix (Solis Biodyne, 08-36-00001-5) on a LightCycler 96 instrument (Roche Diagnostics). Gene expression was normalized to the expression of *18S* and data analyzed and visualized using GraphPad Prism 9. *P values* are indicated in figure legends and were calculated using Two-Way-ANOVA with Tukey’s multiple comparisons test. Primer sequences used in this study are listed in Suppl. Table 1.

### RNA sequencing

#### Library preparation and sequencing

Total RNA was extracted as described in ‘RNA purification’. Samples were cleaned using the RNeasy Mini Kit (Qiagen, 74104) and gDNA removed using the on-column RNase-free DNase set (Qiagen, 79254). RNA quality and concentration were assessed using an Agilent Bioanalyzer device using the RNA 6000 Nano Kit (Agilent, 5067-1511).

Total RNA-Seq libraries were generated from 200 ng of total RNA using Illumina Stranded Total RNA Prep, Ligation with Ribo-Zero Plus kit and IDT for Illumina RNA UD Indexes, Ligation (Illumina, San Diego, USA), according to manufacturer’s instructions. DNA libraries were amplified using 15cycles of PCR. Surplus PCR primers were further removed by two successive purifications using SPRIselect beads (Beckman-Coulter, Villepinte, France). The final libraries were checked for quality and quantified using Bioanalyzer 2100 system (Agilent technologies, Les Ulis, France). Sequencing was performed on the GenomEast platform (IGBMC, Illkirch, France) with 1×50 bp on a NextSeq 2000 System (Illumina).

#### RNA-sequencing analysis

Reads were preprocessed to remove adapter, polyA and low-quality sequences (Phred quality score below 20). After this preprocessing, reads shorter than 40 bases were discarded for further analysis. These preprocessing steps were performed using cutadapt (43) version 4.2. Reads were mapped to rRNA sequences of RefSeq and GenBank databases using bowtie2 (44) version 2.2.8. Reads mapping to rRNA sequences were removed for further analysis. Reads were mapped onto the GRCh38 assembly2 of Homo sapiens genome using STAR (45) version 2.7.10b. Gene expression quantification was performed from uniquely aligned reads using htseq-count (46) version 0.13.5, with annotations from Ensembl version 110 and “union” mode3. Only non-ambiguously assigned reads to a gene have been retained for further analyses.

#### Differential gene expression analysis

RNA-seq raw count data per ENSEMBL ID were analyzed using *DESeq2* (v1.44.0) (47) within the *R* (v4.4.2) programming environment to perform differential gene expression analyses between conditions. A gene was considered differentially expressed with a log2 fold-change (log2FC) cutoff of ±1 and an adjusted p-value (padj) < 0.05 (Benjamini-Hochberg method). To visualize global gene expression changes, volcano plots were generated by plotting the negative log10 adjusted p-values against the log2 fold changes. Within these plots, genes belonging to the BATLAS Brown Adipose Tissue (BAT) (48) signature were specifically highlighted. BAT genes passing both significance thresholds were color-coded to distinguish upregulated and downregulated markers. A lollipop plot was generated to compare the results of the differential gene expression analysis for *UCP1*. The stem height represents the log₂ fold change, and the dot color represents the adjusted p-value.

#### Heatmap of BATLAS genes

The heatmap was generated to visualize the expression patterns of BATLAS genes across all samples. Normalized count data were log2(x+1) transformed to stabilize variance. Row-wise Z-score standardization was applied to center and scale gene expression. Hierarchical clustering of genes (rows) was performed using Euclidean distance as the metric and Ward’s linkage method to minimize within-cluster variance. Samples were ordered by experimental condition (DMSO, Rosi, TAK, TAK+Rosi) to facilitate group comparisons.

#### Single-Sample GSEA (ssGSEA)

To quantify the enrichment of BAT signature within individual samples, single-sample GSEA was performed using ‘gseapy’. Input data consisted of Transcripts Per Million counts, which were log2(x+1) transformed. An enrichment score was calculated for each sample relative to the BATLAS gene set. Statistical differences in ssGSEA enrichment scores between experimental groups were evaluated using a one-way Analysis of Variance.

#### Functional enrichment analyses

The *biomaRt* package (v2.60.1) (49) was used to map ENSEMBL IDs to gene symbols. Gene Ontology (GO) enrichment analysis for GO biological processed (GO BP) was performed using the *clusterProfiler* package (v4.12.6) (50) with differentially expressed genes (up-or down-regulated) as input. To reduce redundancy among enriched GOBP terms, the rrvgo package (v1.18.0) was employed. Semantic similarity matrices were computed based on the org.Hs.eg.db annotation database using the “Rel” (Relevance) metric. Redundant terms were aggregated using the reduceSimMatrix function with a similarity threshold of 0.7; within each cluster, the term with the lowest Benjamini-Hochberg adjusted p-value was selected as the representative. For visualization, the top 15 simplified terms were ranked by significance and displayed as dot plots illustrating the −log10(adjusted p-value) and the number of genes associated with each term.

#### Transcription factor activity analysis

TF activity inference was performed using the *decoupler* (v1.8.0) package in Python (v3.11.7) (51). Log2 fold-change values of upregulated DE genes were used to infer TF activities with the Universal Linear Model method. TF-target interaction networks were obtained from OmniPath (52).

Fig. 6A presents bar plots depicting the inferred most active TFs when comparing TAK-981 + Rosiglitazone versus DMSO control. Fig. 6B-C and Supplementary Fig. 6A show volcano plots of the log2FC and adjusted p-values of the expression of beiging targets of PPARA, PPARG, and SREBP1, respectively.

**Fig. 6:**
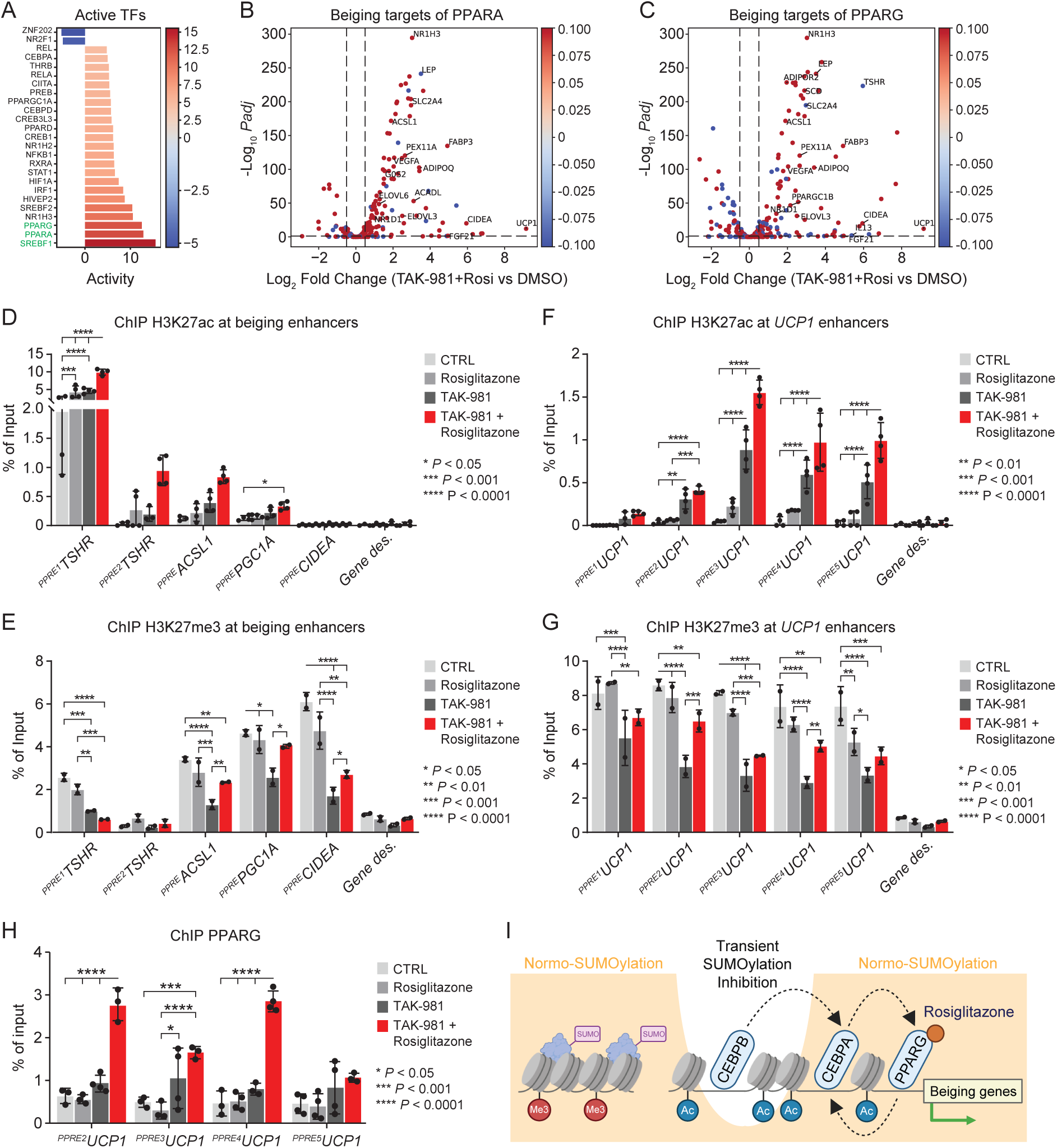
Transient SUMOylation inhibition enhances PPARA/G transactivation and positive epigenetic changes at beiging enhancers. **(A)** *In silico* prediction of mobilized transcription factors based on RNA-seq data, showing the transcription factors mobilized upon treatment with TAK-981 and rosiglitazone, 22 days after adipogenic induction. **(B-C)** Volcano plots showing the PPARA-target genes *(B)* and PPARG-target genes *(C)* activated in response to SUMOylation inhibition and rosiglitazone. Differentially expressed genes were identified by comparing TAK-981 + Rosiglitazone-treated cells to control cells, as described in Fig. 1A and the methods. **(D-E)** ChIP-qPCR assessment of H3K27ac *(D*) and H3K27me3 *(E)* occurrences at selected beiging enhancers, as illustrated in Suppl. Fig 6B-E. Error bars represent the average to the mean of two to four independent experiments. ChIP signals were normalized to inputs, and a gene desert at chromosome 12 (Gene des.) was used as a negative control. Statistical significance was calculated using a Two-Way ANOVA with Tukey’s multiple comparisons test. **(F-G)** ChIP-qPCR assessment of H3K27ac *(F*) and H3K27me3 *(G)* occurrences at *UCP1* enhancers, as illustrated in Suppl. Fig 6F. Error bars represent the average to the mean of two to four independent experiments. ChIP signals were normalized to inputs, and a gene desert at chromosome 12 (Gene des.) was used as a negative control. Statistical significance was calculated using a Two-Way ANOVA with Tukey’s multiple comparisons test. **(H)** ChIP-qPCR assessment of PPARG occurrences at *UCP1* enhancers, as illustrated in Suppl. Fig 6F. Error bars represent the average to the mean of two independent experiments. ChIP signals were normalized to inputs and a gene desert at chromosome 12. Statistical significance was calculated using a Two-Way ANOVA with Tukey’s multiple comparison test. **(I)** Model illustrating the effects of transient SUMOylation inhibition and rosiglitazone treatment on heterochromatin factors, HATs, chromatin structure, and PPARG transactivation during the beige identity specification. Transient SUMOylation inhibition in pre-adipocytes stably primes the chromatin to facilitate the positive effect of Rosiglitazone on the transactivation activity of beiging transcription factors like PPARs. The stable effect of SUMOylation inhibition on the epigenome indicates that SUMOylation in adipoсе stem cells restricts cellular identity shift towards beiging in mature adipocytes.

### ATAC-sequencing

#### Library preparation and sequencing

Samples for Assay for Transposase-Accessible Chromatin sequencing (ATAC-seq) were prepared based on previously published protocols with the Illumina ATAC-seq library preparation kit (20034197) (53–56). Cells were grown and differentiated as described in “Cell culture, differentiation and treatments” as duplicates in 12-well plates. At the day of sample preparation, the cells were trypsinised and 50,000 cells were centrifuged at 500 RCF, 5 min at 4°C. The cells were washed in icecold PBS and centrifuged once more and the supernatant was removed. The cells were resuspended in cold ATAC-seq Lysis Buffer (10mM Tris-HCl, pH 7.5, 10 mM NaCl, 3 mM MgCl_2_ (Sigma, M1028), 0.1% NP-40, 0,1% Tween-20, 0.01% Digitonin (Promega, #G9441)) and incubated on ice for 3 minutes. The nuclei were then washed with the ATAC wash buffer (10mM Tris-HCl, pH 7.5, 10mM NaCl, 3mM MgCl_2_, 01% Tween-20) by inverting 3 times and centrifuged at 500 RCF, 10 min, 4°C. The supernatant was removed and the nuclei pellet was resuspended in 50µL transposition mixture (25µL Tagment DNA Buffer (1x final, Illumina, 20034197), 2.5µL Tagment Tn5 Transposase (100nM final, Illumina, 20034197), 16.5µL 1x PBS, 0.5µL 1% Digitonin (0.01% final), 0.5µL 10% Tween-20 (0.1% final) and incubated at 37°C for 30 min shaking at 1000 rpm. The tagmented DNA was purified with the Zymo DNA Clean and Concentrator-5 Kit (Zymo, D4014) and eluted in 21µL Elution Buffer. The purified DNA fragments were then amplified by PCR to enrich for fragments at 200 – 600 bp length. The PCR reactions were set up with 20µL purified and transposed DNA sample, 2.5µL 25µM Universal primer Ad.1_noMX, 2.5µL 25µM barcoded primer Ad.2 (Ad2.[1-15], Table S1 and (56)) and 25µL NEB Next High-Fidelity 2x PCR Master Mix (NEB, M0541S) and the following thermal cycle settings: 1 cycle: 72°C for 5 min; 1 cycle: 98°C for 30 s; 13 cycles [98°C for 10s, 63°C for 30 s, 72°C for 1 min]; hold at 4°C ∞. The libraries were then purified and size-selected by performing two times a 1.4x bead cleanup with SPRIselect beads (Beckman-Coulter, A63880). In brief, 1.4x the volumes of beads were added to the PCR products and mixed thoroughly before incubating at room temperature for 5 min. The beads with the library were washed twice in 80% fresh ethanol and dried before eluting in 50µL nuclease-free H_2_O. This procedure was repeated a second time with a final elution volume of 25µL. The final concentrations were measured on a Qubit device and the quality of the final size-selected libraries was assessed on a Bioanalyzer with the High Sensitivity DNA kit (Agilent, 76645-152).

#### Library sequencing, alignment, and data processing

Libraries were sequenced on an Illumina NextSeq 2000 sequencer as paired-end 50 base reads. Image analysis and base calling were performed using RTA version 2.7.7 and BCL Convert version 3.8.4. Paired-end sequencing reads were mapped to the Homo sapiens reference genome (assembly hg38) using the ENCODE ATAC-seq pipeline (v2.2.2). Alignment was performed using Bowtie2 (v2.3.4.3) (44) with default parameters, except allowing fragment lengths up to 2 kb (-X2000). Following alignment, SAMtools (v1.9) (57) was used to filter reads (-F 1804-q 30); unmapped reads, mate unmapped reads, non-primary alignments, reads failing platform quality checks, and low-quality reads (MAPQ < 30) were removed. PCR duplicates were marked using Picard MarkDuplicates (v2.20.7) and subsequently removed using SAMtools (-F 1804). For visualization, BigWig files were generated using deepTools bamCoverage (v3.5) (58). Coverage was calculated in 10 bp bins (-bs 10), normalized to Counts Per Million (CPM), with reads extended and non-covered regions skipped. An effective genome size of 2,913,022,398 was used.

#### Chromatin accessibility peak calling and annotation

Peak calling was performed on individual samples using MACS2 (v2.2.4) (59) via the ENCODE ATAC-seq pipeline. The following parameters were used to account for the Tn5 transposase shift and fragment size: --shift 75 --extsize 150 --nomodel-B --SPMR --keep-dup all --call-summits. The “optimal overlap” peak set was retained for downstream analyses. Peaks were annotated relative to genomic features using HOMER (v4.11.1) (60) with annotations extracted from Ensembl (v110).

#### Differential accessibility analysis and temporal clustering

To perform differential analysis, a union set of all detected peaks (Open Chromatin Regions, OCRs) was generated using Bedtools merge (v2.30.0) (61). Read counts for each merged peak were computed using Bedtools intersect (v2.30.0) and normalized using the method described in (62) as implemented in DESeq2 (v1.34.0) (47). Differential binding was assessed using the Wald test in DESeq2. P-values were adjusted for multiple testing using the Benjamini-Hochberg procedure, with significance defined as an adjusted p-value < 0.05 and a |log2 Fold-Change| > 0. For time-series analysis (DMSO/TAK), significant changes were identified using a Likelihood Ratio Test (LRT) with an adjusted p-value ≤ 0.05. These differentially bound regions were clustered based on their binding profiles using the fuzzy c-means algorithm implemented in the Mfuzz package (v2.62.0) (63).

#### Functional enrichment analysis of differentially accessible regions

Genomic regions within each temporal cluster were merged using the ‘reducè function from the GenomicRanges package to identify distinct features; clusters with fewer than 20 regions were excluded. Functional enrichment analysis was performed using the ChIP-Enrich package (64) with the Homo sapiens reference genome (hg38). Enrichment was tested against Gene Ontology Biological Processes (GOBP) using the ‘nearest_tss’ locus definition, assigning peaks to the gene with the closest Transcription Start Site. To generate the heatmap, the top 15 significant terms (FDR < 0.05) from each cluster, ranked by FDR and Odds Ratio, were aggregated to define the universe of terms. The enrichment values (Odds Ratios, OR) were log2-transformed log2(OR+1) and visualized using the ‘pheatmap’ package, with hierarchical clustering applied to both rows and columns. The size the of individual points on the dotplot depend on the OR value.

#### Genome-wide differential Transcription Factor activity

Genome-wide differential transcription factor activity (Fig. 5D-F) was quantified using diffTF (version 1.9) (65) in basic mode. Transcription Factor Binding Sites obtained from the UniBind database (66) were intersected with the consensus ATAC-seq peaks, applying a 100 bp extension in both directions. For each TFBS, accessibility fold-changes were calculated using the design formula ∼ conditionSummary and stratified into 10 bins based on GC content. Differential activity was estimated by comparing the fold-change distribution of target sites against a background of TFBS with similar GC content using the analytical approach. Variance was estimated using 1,000 bootstrap replicates. TFs were considered differentially active if they met a significance threshold of Benjamini-Hochberg adjusted FDR < 0.01 and TF Activity Change > |0.05|.

To visualize global TF activity changes, volcano plots were generated by plotting the negative log10 adjusted p-values against the differential binding scores computed by diffTF. All TFs with an absolute differential binding score > 0.05 and pAdj < 0.001 were colored depending on the sign of the differential binding score (negative: blue and positive: red). The global temporal evolution of transcription factor activities was represented as a connected dot plot, with time points on the horizontal axis and differential binding score on the vertical axis. Transcription factors of interest are highlighted, and dot size represents −log₁₀(adjusted p-value).

Transcription factor activity was represented using density plots of the log₂ fold change in activity at each transcription factor binding site (TFBS) for CEBP, NFIL3, and PPARG (blue). The density of the log₂ fold change in activity at each background TFBS was also shown on each plot (Suppl. Fig 5C-D).

#### Thermogenesis pathway visualization

The “Thermogenesis” KEGG pathway was visualized with differentially expressed genes between TAK-981 + Rosiglitazone versus DMSO using the *pathview* (v1.44.0) R package (67) (Suppl. Fig. 4A).

#### Protein extraction and western blotting

For UCP1 and TBP detection (Fig. 2E), pCREB/pATF1 and TBP detection after treatment with Rp-8-CPT-cAMPS(Fig. 4F), and PKA phospho-substrates and TBP detection by Western Blot (Fig. 4B), cells were washed in ice-cold PBS prior lysis in ice-cold RIPA buffer (50mM Tris-HCl pH 7.5, 150mM NaCl, 5mM EDTA, 1% NP-40, 0.5% Sodium deoxycholate, 0.1% SDS), freshly supplied with protease inhibitor (Roche) and phosphatase inhibitor (Roche). Lysates were centrifuged at 13,000 rpm for 10 minutes at 4°C. To eliminate lipid carryover, the supernatant was filtered through a RNeasy column (Qiagen, 74104) and centrifuged at 10,000 rpm for 1 min at 4°C. The flow through was collected and protein concentrations were measured with the Braford Assay. 20 - 40µg of protein were used for SDS-PAGE and proteins of interest detected by Western Blots.

For SUMO2/3 (Fig. 1B), p38, p-p38, p-ATF2, p-ATF1, p-CREB (Fig. 4D) and CEBPB, PPARG and TBP (Fig. 5H, I) Western Blots, cells were lysed in 2x Laemmli Buffer and sonicated for 2 cycles (30 sec on/30 sec off). Lysates were centrifuged at 13,000 rpm for 10 minutes at 4°C. To eliminate lipid carryover, the supernatant was filtered through a RNeasy column (Qiagen, 74104) and centrifuged at 10,000 rpm for 1 min at 4°C. Protein concentrations were determined with the Bradford Assay. 15 – 50µg of protein were used for SDS-PAGE and proteins of interested detected by Western Blots.

For UCP1 and γ-tubulin detection in differentiated hTERT A41hWAT-SVF at Day 21 after pausing Rosiglitazone administration by Western Blot (Fig. 2I), the cells were washed in ice-cold PBS prior to lysis with 2x Laemmli Buffer. The cells were boiled for 5 min at 95 °C and the lysate were passed through a 26g needle 3 to 5 times before centrifuging for 10min at 13,000 rpm. Equal amounts were loaded on a SDS-PAGE based on the ratio of γ-tubulin as loading control.

For UCP1, γ-tubulin, PPARG1/2 and TBP detection in hASCs by Western Blot (Fig. 2F, 5J, Supp. Fig. 4D), the cells were washed in ice-cold PBS prior to lysis with adipocyte lysis buffer (50 mM Tris pH 7.4, 0.27 M sucrose, 1 mM Na-orthovanadate pH 10, 1 mM ethylenediaminetetraacetic acid (EDTA), 1 mM ethylene glycol-bis(β-aminoethyl ether)-*N,N,N*’,*N*’-tetraacetic acid (EGTA), 10 mM Na β-glycerophosphate, 50 mM NaF, 5 mM Na pyrophosphate, 1% (w/v) Triton X-100, 0.1% (v/v) 2-mercaptoethanol and cOmplete™ Protease Inhibitor Cocktail, (Roche Diagnostics, Meylan, France). The lysates were passed through a 26g needle 3 to 5 times and centrifuged at 13,000rpm for 10 min at 4°C. Protein concentrations were determined with the Bradford assay and 30µg protein were used for SDS-PAGE and Western Blot.

The following antibodies were used: UCP1, abcam, ab209483; PKA phospho-substrates, Cell Signaling 9624; p-CREB / p-ATF1, Cell Signaling 9198; p38 MAPK, Cell Signaling 9212; p-p38 MAPK, Cell Signaling 9211; p-ATF2, Cell Signaling 24329; SUMO2/3, abcam, ab81371; PPARG, Cell signaling 2443 and TBP, Protein Tech, 22006-1-AP or abcam, 282715; γ-tubulin, Sigma, T5326: CEBPB, Santa Cruz, sc-7962 Quantifications were performed using FiJi (68).

#### Mitochondrial respiration analysis

Human adipocytes were differentiated as described in ‘Cell culture, differentiation and treatment’ in 24 multiwell plates (Seahorse). Oxygen Consumption Rate (OCR) and Extracellular Acidification Rate (ECAR) were measured using the XF Cell Mito Stress Test on a XF24 extracellular Flux Analyzer (Seahorse Agilent). During the assay, 2µM Oligomycin (O4876, Sigma) and 2.5µM BAM15 (HY-110284, MedChemExpress) were added as indicated. For the quantification of basal, maximal and uncoupled respiration as well as proton leak, the data were normalized to cell number as assessed by crystal violet staining.

#### Crystal violet staining

Crystal Violet staining was performed as in (69). Cells were washed twice in PBS and fixed in 4% formaldehyde for 10 minutes at room temperature. Afterwards the cells were washed in PBS and incubated in Crystal violet staining (Ås Produksjonslab AS, 0.1% working concentration in PBS) for 4 – 7 minutes at room temperature. Unbound dye was washed off at least 5 times with mqH_2_O, followed by solubilizing the dye with 33% acetic acid for 5 minutes rocking. Finally, the OD was measured at 550nm.

#### Chromatin immunoprecipitation (ChIP)

Human white adipocytes were differentiated as described in ‘Cell culture, differentiation and treatment’ in 10cm-dishes. Cells were washed twice in PBS and collected in Digitonin lysis buffer with sodium-butyrate (10mM Tris-HCl, pH 7.4, 10mM NaCl, 3mM MgCl_2_, 20mM Sodium-butyrate, 0,01% Digitonin, 0.1% Tween-20, 0.1% NP-40) (70). To obtain nuclei, the cells were dounced (35 strokes) using pestle B. Cell lysis was assessed and confirmed with a phase-contrast microscope. Nuclei were centrifuged at 500g for 10 min at 4°C. The nuclei pellet was washed twice in Digitonin lysis buffer and resuspended in nuclear pellet buffer NPB (10mM HEPES, 1.5 MgCl_2_, 250mM sucrose, 0.1% NP-40, protease inhibitor cocktail, 1mM PMSF and 20mM Sodium-butyrate in PBS). Chromatin was crosslinked with 1% Formaldehyde for 8 min. Formaldehyde was neutralized using 125mM Glycine for 10min. After two washes with NPB, the nuclei were resuspended in nuclear lysis buffer (50 mM Tris-HCl pH 7.5, 1% SDS, 10 mM EDTA, 20 mM N-ethyl maleimide and protease inhibitor cocktail) and incubated at 4°C for 2 h. Lysates were sonicated for 10 cycles (30 s on/30 s off) at 4°C using a Bioruptor Pico sonicator (Diagenode). After sonication, lysates were centrifuged at 14,000 rpm for 10 min at 4°C. Protein concentration was assessed using the Bradford assay and 30 µg protein were used for histone immunoprecipitation and 250µg for PPARG immunoprecipitation. Input samples (10 µg) were saved. Samples were diluted 10-fold in the immunoprecipitation buffer (1.1% Triton X100, 50 mM Tris-HCl, pH 7.5, 167 mM NaCl, 5 mM N-ethyl maleimide, 1 mM EDTA, 0.01% SDS and protease inhibitor cocktail). Immunoprecipitations were carried out with antibodies for H3K27ac (ab4729, abcam), H3K27me3 (C15410195, Diagenode), PPARG (2443S, Cell Signaling and C15410367, Diagenode) and Dynabeads Protein A or G (10002D, 10004D, ThermoFisher Scientific) overnight at 4°C. Beads were then washed 2 times in low-salt buffer (50 mM Tris-HCl, pH 7.5, 150 mM NaCl, 1% Triton X100, 0.1% SDS, 1 mM EDTA), 2 times in high-salt buffer (50 mM Tris-HCl, pH 7.5, 500 mM NaCl, 1% Triton X100, 0.1% SDS, 1 mM EDTA) and in TE buffer (10 mM Tris-HCl, pH 7.5, 0.2% Tween-20, 1 mM EDTA). Elution was done two time in 50µl of 100 mM NaHCO3, 1%SDS at 65◦C for 10 min under agitation. Chromatin crosslinking was reversed at 65°C for 5 h with 280 mM NaCl and 0.2 µg/ml RNase DNase free (11119915001, Roche). Proteins were then digested using 0.2 µg/ml of Proteinase K (03115828001, Roche) during 1 h at 65°C. DNA from immunoprecipitations and inputs were purified using the Qiagen PCR purification kit. The resulting samples were further used for qPCR as described before.

#### Genome tracks

The UCSC genome browser (71) (hg19) was used to generate Suppl. Fig. 6G. The GENCODE V47lift37 (72), NCBI RefSeq (73), and HGNC (74) tracks provide information about gene annotations. The JASPAR CORE 2024 track (75) was filtered to provide transcription factor binding site (TFBS) predicted with JASPAR profiles associated with Ppara (MA2338.1), PPARA::RXRA (MA1148.2), PPARG (MA0066.2), and Pparg::Rxra (MA0065.3). The PPRE_hg19.bed track provides the localization of the TFBSs considered to design the amplicons, which were lifted over from hg38 to hg19. The tracks PPARGbriteRep1 and PPARGbriteRep2 represents bigWig of two PPARG ChIP-seq replicate experiments in mature human adipose-derived stem cells (76) and were obtained from GEO (GSM1443810 and GSM1443812). The tracks H3K27acBriteRep1 and H3K27acBriteRep2 represents bigWig of two H3K27ac ChIP-seq replicate experiments from the same cell types (76) and were obtained from GEO (GSM1443806 and GSM1443808).

## Results

### Transient SUMOylation inhibition in human pre-adipocytes stably steers the transcriptome towards beiging in a rosiglitazone-dependent manner

To investigate the role of SUMOylation in adipocyte beiging, we transiently treated human immortalized hTERT A41hWAT-SVF pre-adipocytes (77) with TAK-981, a selective inhibitor of the SUMOylation pathway (78,79), for 48h prior to adipogenic induction (Fig. 1A). Following adipogenic induction, cells were treated with rosiglitazone, a PPARG agonist and well-established inducer of adipocyte beiging (80). This rosiglitazone treatment is a prolongation of the required 7 days treatment necessary to achieve white adipocyte differentiation in this cellular model without triggering beiging (37) and will be referred to as “+ Rosi” for the sake of clarity. Finally, a third experimental condition combined TAK-981 + Rosi to assess potential cumulative effects (Fig. 1A).

Western blot analysis for SUMO-2/3 confirmed the transient nature of SUMOylation inhibition. SUMO conjugates levels dropped at day 0 and recovered a few days following transient TAK-981 treatment (Fig. 1B). Conversely, free SUMO accumulated immediately after the treatment and slowly decreased as SUMOylation recovered (Fig. 1B). It is worth mentioning that rosiglitazone did not have a noticeable effect on global SUMOylation levels (Suppl. Fig. 1A). To evaluate the impact of these treatments on adipocyte differentiation, we performed Oil-Red-O staining after SUMOylation recovery at day 22. As expected, rosiglitazone treatment enhanced lipid accumulation compared to control cells (Suppl. Fig. 1B) (80). Notably, an increase in lipid accumulation was observed following TAK-981 treatment, suggesting that SUMOylation inhibition in white pre-adipocytes promotes adipogenesis, consistent with our previous findings in mouse beige pre-adipocytes (81). However, the combination of TAK-981 and rosiglitazone did not enhance adipogenesis beyond the effects observed with TAK-981 only (Suppl. Fig. 1B).

To assess the impact of transient SUMOylation inhibition in pre-adipocytes on adipocyte fate in an unbiased manner, we performed RNA sequencing (RNA-seq) at day 21 in cells subjected to the treatments described in Fig. 1A. Differentially expressed gene (DEG) analysis (log₂FC > 1 or log₂FC < −1; padj < 0.05) revealed significant transcriptome alterations in all three conditions compared to control cells (Suppl. Table 2). TAK-981 alone led to 336 downregulated and 1,169 upregulated genes (Fig. 1C and Suppl. Table 2), which was consistent with previous reports indicating that SUMOylation functions as both a facilitator or repressor of gene expression, with a predominant role in gene repression (5,82–86). Rosiglitazone alone resulted in 524 downregulated and 960 upregulated genes (Fig. 1D Suppl. and Table 2). Co-treatment with TAK-981 and rosiglitazone induced the most extensive transcriptomic changes, with 953 downregulated and 2,064 upregulated genes (Fig. 1E Suppl. and Table 2). We also compared TAK-981 + rosiglitazone to TAK-981 alone and identified 652 upregulated and 447 downregulated genes (Suppl. Fig. 1C and Suppl. Table 2). We finally compared TAK-981 + rosiglitazone to rosiglitazone alone. Consistent with the repressive role of SUMOylation, we identified 682 upregulated and 190 downregulated genes (Suppl. Fig. 1D and Suppl. Table 2).

Gene ontology analysis of biological processes (GO BP) (87) demonstrated a strong enrichment for fatty acid and lipid metabolic pathways among the upregulated genes across all five comparisons (Suppl. Fig. 2A-E, left and Suppl. Table 3), consistent with the established role of lipid metabolism in both white and beige adipocytes (88). More notably, the analysis revealed significant enrichment for processes related to brown and beige adipocytes, including “positive regulation of cold-induced thermogenesis,” “temperature homeostasis,” “adaptive thermogenesis,” and “brown fat cell differentiation.” These GO terms were further enriched in co-treated cells compared to those receiving individual treatments (Suppl. Fig. 2A-E, left and Suppl. Table 3). Interestingly, when using rosiglitazone as a baseline to specifically identify pathways whose regulation was enhanced by SUMOylation inhibition, we observed the strongest enrichment for brown and beige adipocyte-related processes across all comparisons (Suppl. Fig. 2E, left and Suppl. Table 3). This suggests that, while rosiglitazone alone induced the expected beiging genes (Suppl. Fig. 2B), SUMOylation inhibition in pre-adipocytes acted as a booster for pathways that promote beiging in response to rosiglitazone.

Genes down-regulated by TAK-981 alone or in combination with rosiglitazone are enriched for BPs like differentiation and development of numerous cell types, including ossification (Suppl. Fig. 2A-E, right and Suppl. Table 3), which is consistent with the effects of SUMOylation inhibition in other systems and adipocytes (81). However, despite significant transcriptional alterations, these comparisons did not result in the enrichment of beiging-specific GO BPs, suggesting that SUMOylation inhibition in pre-adipocytes selectively reinforces beiging in rosiglitazone-treated cells.

To further characterize the effects of TAK-981 on genes associated with beiging, we utilized BATLAS, a recently published gene expression signature for brown adipose tissue (BAT) (48). A heatmap was generated to visualize the expression patterns of BAT genes across all samples. Both TAK-981 and rosiglitazone administered individually increased the expression of most BATLAS genes compared to control cells (Fig. 1F). Strikingly, the upregulation of BATLAS genes was more pronounced in cells receiving co-treatment with TAK-981 and rosiglitazone (Fig. 1C-E, Fig. 1F and Suppl. Fig. 1C,D). Importantly, the expression of the key beiging marker *UCP1* increased from single treatments to the combination treatment with TAK-981 and rosiglitazone (Fig. 1C-G). These results suggest a transcriptional shift towards a beiging phenotype in cells treated with TAK-981 and rosiglitazone.

To further validate this finding, we performed single-sample gene set enrichment analysis (ssGSEA) using BATLAS genes as a reference [as described in (40)]. As anticipated, this analysis indicated a significant shift in gene expression towards a BAT profile consistent with beiging in cells treated with rosiglitazone (Fig. 1H). More unexpectedly, while *UCP1* was not significantly upregulated (see Fig. 1G), treatment with TAK-981 alone also elicited a comparable shift due to the upregulation of numerous other BAT genes (Fig. 1H). Finally, the combination of TAK-981 and rosiglitazone showed the strongest correlation between our expression data and BATLAS genes (Fig. 1H).

Collectively, these findings suggest that SUMOylation in pre-adipocytes acts as a barrier to inhibit transcriptional reprogramming towards a beige adipocyte phenotype induced by rosiglitazone.

### Transient SUMOylation inhibition primes pre-adipocytes to stably express UCP1

We next sought to validate RNA-seq data using RT-qPCR. We first examined the effects of these treatments on the expression of adipogenic genes. Both rosiglitazone and TAK-981 upregulated the beige-specific gene *PPARG2* compared to control cells (Fig. 2A). Co-treatment with both molecules slightly enhanced *PPARG2* expression beyond the levels observed with either treatment alone (Fig. 2A). A similar trend or an additive effect was observed for other adipogenic markers (Suppl. Fig. 3A). This is consistent with RNA-seq data (Suppl. Table 2) and with the lipid accumulation observed in Suppl. Figure 1B. Such an enhanced adipogenic transcription has been shown to cooccur with beiging, notably for PPARG, which is a transactivator of *UCP1* gene (89,90).

Next, we quantified the expression of *UCP1*, a hallmark of beige adipocyte conversion and a key mediator of mitochondrial uncoupling (21). While *UCP1* expression was undetectable in control cells and weakly induced by either rosiglitazone or TAK-981 alone, co-treatment with both molecules led to a strong increase in *UCP1* expression (Fig. 2B). Stinkingly, the expression of *UCP1* remained strong 32 days after transient TAK-981 treatment suggesting an imprinting effect of SUMOylation inhibition leading to constitutive *UCP1* expression, which differs from other SUMO-regulated genes, such as the *IFN1B* in myeloid cells (91).

Next, we sought to validate these data in primary human adipose stem cells (hASC) from healthy donors (39–41). These cells presented a fat accumulation pattern comparable to immortalized pre-adipocytes (Suppl. Fig. 1B and Suppl. Fig. 3B). Furthermore, as in immortalized pre-adipocytes (Fig. 2A), we observed that rosiglitazone and TAK-981 significantly upregulated *PPARG* expression compared to control cells (Fig. 2C). However, co-treatment with both molecules further enhanced *PPARG* expression beyond the levels observed with either treatment alone (Fig. 2C). A similar trend was observed for the other adipogenic genes tested previously in immortalized pre-adipocytes (Suppl. Fig. 3C). Finally, we also observed a significant, additive increase of *UCP1* mRNA levels in co-treated hASCs compared to single treatments (Fig. 2D). These data indicate that co-treatment with TAK-981 and rosiglitazone significantly increased the expression of *PPARG* and *UCP1*, beyond individual treatments, in primary hASCs.

To determine whether the observed increase in *UCP1* mRNA levels correlated with higher UCP1 protein expression, we performed western blot analysis in pre-adipocytes using differentiated brown adipocytes (hTERT A41hBAT-SVF, see methods) as a positive control (77). Consistent with *UCP1* transcript levels in RNA-seq and RT-qPCRs (Fig. 1C,E and Fig. 2B and Suppl. Table 2) and previous studies (80), UCP1 protein expression was only weakly detectable at day 32 in cells treated with rosiglitazone alone (Fig. 2E). A faint UCP1 band was also observed in TAK-981-treated cells at day 32 (Fig. 2E). In contrast, co-treatment with TAK-981 and rosiglitazone resulted in a robust increase in UCP1 protein levels, which became evident as early as day 22 and which reflected the dynamics of *UCP1* mRNA levels (Fig. 2B,E and Suppl. Table 2). We validated these data in primary hASCs, where UCP1 protein levels were stronger in cells co-treated with TAK-981 and rosiglitazone than in the other conditions, which correlated with the pattern of *UCP1* mRNA dynamics across conditions in these cells (Fig. 2D,F).

Next, we asked whether transiently inhibiting SUMOylation in pre-adipocytes is sufficient to stably induce a beiging fate. We also wanted to confirm that the PPARG agonist rosiglitazone acts downstream to finalize the transition from white to beige fat cells. To test this, we interrupted the rosiglitazone treatment during six days and measured *UCP1* expression (Fig. 2G, lower part). If beiging is successfully imprinted by TAK-981, *UCP1* levels should increase even with the delayed rosiglitazone treatment. However, if the beiging fate is not imprinted, the cells will revert to their original white fate, and rosiglitazone will have little effect on *UCP1* expression. As shown in Fig. 2B, under continuous rosiglitazone treatment, combining rosiglitazone with TAK-981 significantly increased *UCP1* expression compared to either treatment alone (Fig. 2H, see Continuous). Remarkably, even when rosiglitazone treatment was interrupted six days, Rosiglitazone-induced *UCP1* expression remained significantly stronger in cells pre-treated with TAK-981, whereas rosiglitazone alone had little effect (Fig. 2H, see Gap). Interestingly, the combination of TAK-981 and delayed rosiglitazone resulted in even higher *UCP1* expression than during continuous treatment. This suggests that a form of “rosiglitazone memory” is established after culturing in the induction medium from days 0 to 7, allowing for a potent response when combined with TAK-981. Since rosiglitazone is an agonist of PPARG involved both in adipogenesis and beiging, we analysed *PPARG* expression and adipogenesis under the same conditions. In continuous conditions of rosiglitazone treatment *PPARG* expression was comparable to Fig. 2A and introducing a six-day gap had no effect on *PPARG* mRNA levels or adipogenesis (Suppl. Fig. 3D,E).

Finally, we measured UCP1 protein levels under these conditions. As shown in Fig. 2E, UCP1 protein levels were much higher when combining continuous rosiglitazone treatment with TAK-981 compared to either treatment alone (Fig. 2I). Similarly, even when rosiglitazone treatment was delayed by six days, the combination with TAK-981 still strongly increased UCP1 protein levels (Fig. 2I). However, despite observing higher *UCP1* mRNA levels with the six-day gap (Fig. 2H), the protein levels remained like those seen in continuous treatment conditions, indicating a possible saturation effect (Fig. 2I).

Together, these data show that transient inhibition of SUMOylation in human pre-adipocytes before adipogenic induction primes the cells to respond more quickly and strongly to rosiglitazone, effectively enhancing the expression of beiging genes, including the key beiging factor UCP1.

### *UCP1* derepression leads to mitochondrial uncoupling

Beige adipocytesare characterized by the expression of thermogenic genes such as UCP1, whose activation drives mitochondrial uncoupling, increased oxidative metabolism and energy expenditure. Measuring Extracellular Acidification Rate (ECAR) provides an indirect readout of UCP1 function because UCP1 activation in brown and beige adipocytes enhances glycolysis and proton leak-driven respiration, both of which contribute to extracellular acidification. In addition, Oxygen Consumption Rate (OCR) reflects increased mitochondrial respiration, confirming UCP1 activation.

Thus, we measured OCR and ECAR at day 22 in cells subjected to the treatments described in Fig. 1A, using Seahorse analysis with differentiated brown adipocytes as a positive control (77). Consistent with UCP1 upregulation, pre-treatment with TAK-981 and then rosiglitazone resulted in a significant increase in OCR and ECAR, including both basal and maximal respiration (Fig. 3A-C). Finally, integrating OCR and ECAR data at the peak of mitochondrial activity, i.e. right before Rotenone/Antimycin A injection, revealed a pronounced shift toward a highly energetic metabolic state in TAK-981 and rosiglitazone-treated cells, approaching the metabolic profile of brown adipocytes (Fig. 3D). These metabolic changes are consistent with beiging in human adipocytes (76) and indicate that transient SUMOylation inhibition is sufficient to change the fate of pre-adipocytes from differentiating into energy-storing white adipocytes to energy-dissipating beige adipocytes in a rosiglitazone-dependent manner.

These data, together with those in Fig. 1 and 2, show that SUMOylation in pre-adipocytes limits beiging capacity.

### Transient inhibition of SUMOylation in pre-adipocytes stably enhances cAMP-PKA-p38 signaling to facilitate beiging

To investigate the molecular basis of the transcriptional and metabolic remodeling observed above, we subjected our RNA-seq data to an unbiased KEGG pathway enrichment analysis. It revealed that TAK-981 + rosiglitazone treatment, compared to DMSO control, resulted in significant upregulation of genes involved in the pathway “adaptive thermogenesis” including components of β3-adrenergic receptor (β3-AR)-PKA-p38 signaling, a key pathway in beiging (Suppl. Fig. 4A). During adaptive thermogenesis, this pathway is initiated by β3-AR activation, leading to PKA-dependent phosphorylation of nuclear substrates (e.g., CREB, JMJD1A) that drive beiging gene expression (e.g., *PPARGC1A*, *PPARG, UCP1*) (88). PKA also phosphorylates cytoplasmic targets such as PLIN1 and HSL, promoting lipolysis and activating p38, which in turn phosphorylates ATF2 to induce *UCP1* expression (Suppl. Fig. 4A) (92,93). We thus hypothesized that SUMOylation blunts the expression of genes involved in the cAMP-PKA-p38 pathway during adaptive thermogenesis (Suppl. Fig. 4B).

Specific analysis of DEGs from Fig. 1 confirmed overexpression of genes such as *Gs*, adenylate cyclase subunits, *JMJD1A*, *PPARGC1A*, and *PPARG* (Fig. 4A and Suppl. Fig. 4A). In contrast, we observed reduced expression of *CB1R*, a known adenylate cyclase inhibitor, suggesting further contribution of SUMOylation inhibition to sustained PKA activation (Fig. 4A and Suppl. Fig. 4A). Furthermore, downstream genes involved in thermogenesis, such as lipolysis-associated genes (*PLIN1*, *PNPLA2*, *LIPE*) were also upregulated, further supporting beiging activation. Finally, genes encoding OXPHOS components were also significantly upregulated, correlating with the high-energy metabolic state of TAK-981 + rosiglitazone-treated cells (Fig. 4A and Suppl. Fig. 4A). These data indicate that inhibition of SUMOylation supports cAMP-PKA-p38 signaling by enhancing pathway genes expression.

Next, to investigate whether these transcriptional changes translated into enhanced pathway activity, we examined the phosphorylation of PKA and p38 substrates (Suppl. Fig. 4C). Western blot analysis using a pan-PKA phospho-specific substrate antibody showed that TAK-981 alone was a more potent activator of PKA than rosiglitazone for certain targets (Fig. 4B). Combining the two drugs yielded a more significant activation of PKA than observed with Rosiglitazone alone but not with TAK-981 alone (Fig. 4B and C). Specific analysis of CREB phosphorylation, a direct PKA target, confirmed that both TAK-981 and rosiglitazone enhanced CREB phosphorylation, while their combination yielded an additional increase (Fig. 4D and E). Notably, ATF-1 phosphorylation followed a similar pattern (Fig. 4D). Importantly, treatment with the PKA inhibitor RP-8-CPT-cAMPS strongly suppressed the phosphorylation of both CREB and ATF1 induced by SUMOylation inhibition, further confirming enhanced PKA signaling under these conditions (Fig. 4F). Furthermore, pharmacological activation of cAMP-PKA signaling in hASCs with forskolin (FSK), led to increased CREB and ATF1 phosphorylation and UCP1 expression (Suppl. Fig. 4D). This is consistent with previous studies indicating the link between cAMP-PKA signaling and beiging (94,95). Finally, the phosphorylation of p38 was also significantly increased by both TAK-981 and rosiglitazone, although there was no additional effect of their combination (Fig. 4D and G). However, phosphorylation of p38’s thermogenic target, ATF2, was significantly stronger when both drugs were combined (Fig. 4D and H).

Together, these findings suggest that SUMOylation restricts beiging via two non-exclusive mechanisms: 1) Transcriptional repression of key components in the β3-AR-cAMP-PKA-p38 pathway, as well as genes encoding downstream OXPHOS proteins; 2) Direct negative regulation of PKA and p38 kinase activities, limiting their capacity to drive thermogenic gene expression.

### Transient SUMOylation inhibition in pre-adipocytes triggers stable remodeling of chromatin accessibility and dynamic mobilization of CEBPs and PPARG

Next, we sought to get mechanistic insight into how transient SUMOylation inhibition in pre-adipocytes primes the stable expression of beiging genes, including *UCP1,* in mature adipocytes. Given the documented increase in chromatin accessibility observed in differentiated beige adipocytes compared to white adipocytes (41) and strong involvement of SUMOylation in restraining chromatin accessibility (96), we hypothesized that TAK-981 induced chromatin opening at key enhancers involved in adipogenic and beiging transcription. To address acute effect of TAK-981 on the chromatin, we thus performed ATAC-seq experiments immediately after the end of TAK-981 and DMSO (CTRL) treatments, in pre-adipocytes, before adipogenic induction (D0) as well as 12 hours (12h) and 7 days (D7) after adipogenic induction (Fig. 5A). Note that, unlike in previous experiments, these two conditions were sufficient to address the question since rosiglitazone is required to initiate differentiation (Fig. 1A). This setup thus also allowed us to primarily focus on TAK-981 effects using rosiglitazone as a baseline.

We first performed a time series analysis of significant ATAC-seq peaks combined with clustering, which showed pronounced changes in chromatin accessibility over time in TAK-981-treated cells compared to CTRL cells (Fig. 5B). Notably, clusters 4, 8 and 10 featured enhanced accessibility at D7 while clusters 1, 2, 3, 5, 7 and 9 displayed reduced accessibility. This indicates the establishment of a novel chromatin accessibility landscape seven days after adipogenic induction. These clusters were linked to genes involved in biological processes such as cell morphogenesis, cell migration / mobility or nucleosome assembly for the most prominent (Suppl. Fig. 5A,B and Suppl. Table 4). Interestingly, cluster 6 featured increased accessibility immediately after the treatment (Fig. 5B) and was the sole cluster to include loci involved in “brown fat cell differentiation” (Fig. 5C; Suppl. Fig. 5A,B; Suppl. Table 4). Importantly, clusters 6 and 2 also shared the term “adaptive thermogenesis” (Fig. 5C; Suppl. Fig. A,B; Suppl. Table 4). These data indicate that TAK-981 induced a large remodeling of chromatin accessibility, including loci involved in beiging transcription, as early as day 0.

Next, we wanted to identify the TFs affected by transient SUMOylation inhibition. We thus performed a footprint analysis of ATAC-seq data to reveal up-and down-regulated differentially bound TFs (dbTF; Differential binding score > 0.05 and *pAdj* < 0.001). At D0, we identified 31 downregulated and 35 upregulated dbTFs (Fig. 5D and Suppl. Table 5). Amongst the most significantly upregulated dbTFs, we found the adipogenic factors CEBPs and NFIL3, which fits with a beiging signature we recently characterized in hASCs (41). At 12h after adipogenic induction, we identified 29 downregulated and 54 upregulated dbTFs (Fig. 5E and Suppl. Table 5). Also fitting with a beiging signature, we observed the downregulation of AP-1 (FOSL2, FOS, JUND, JUNB) while CEBPs and NFIL3 footprints remained strongly upregulated (Fig. 5E,G and Suppl. Fig. 5C and Suppl. Table 5). Finally, at D7 after adipogenic induction, we identified 123 downregulated and 41 upregulated dbTFs (Fig. 5F and Suppl. Table 5). At this point, NFIL3 footprints decreased (Fig. 5F and Suppl. Fig. 5C) while CEBPs remained highly mobilized (Fig. 5F,G and Suppl. Fig. 5C). This includes CEBPA, a known transactivator of *PPARG* (Fig. 5G). In parallel, PPARG became highly significantly mobilized, which correlates with its function as main activator of *UCP1* expression (Fig. 5F,G and Suppl. Fig. 5D and Suppl. Table 5).

Given the increase in CEBPs footprints at D0, we hypothesized that TAK-981 had a direct effect on the SUMOylation status and the activity of these proteins. To address this, we analyzed the adipogenic SUMO target CEBPB (5,12) by western blots at D0. These experiments revealed the disappearance of a previously identified SUMO-CEBPB form (12) upon TAK-981 treatment when compared to control cells (Fig. 5H). Furthermore, consistent with the function of CEBPB SUMOylation, which is to facilitate proteasome-mediated degradation (12), CEBPB protein levels increased at D0 but also at D22 (Fig. 5H,I). In parallel, PPARG protein levels also increased in TAK-981 treated pre-adipocytes and hASCs at the end of the time course (Fig. 5I,J). Importantly, in both cell types, we observed the enrichment of the beige-specific PPARG2 isoform (40), especially in co-treated cells, which further supports the reprogramming toward a beige identity.

Together, these data show an immediate remodeling of chromatin accessibility in response to TAK-981, which is characterized by the mobilization of adipogenic pioneer factors CEBPs. Then, a stable alternative accessibility landscape is established, featuring the strong mobilization of CEBPA and PPARG. Consistently, TAK-981 stabilized CEBPB and PPARG proteins throughout differentiation, which is coherent with increase mRNA levels detected in Fig. 2 and supports the activity of beiging enhancers and genes.

### Transient SUMOylation inhibition in pre-adipocytes initiates a stable epigenetic reprogramming of PPARA/G enhancers to drive beiging in mature adipocytes

Given that previous studies linked CEBPB and CEBPA to the transactivation activity of PPARG (97), our ATAC-seq experiments suggested that the early mobilization of CEBPs upon TAK-981 treatment led to the activation of PPARG to sustain beiging genes expression downstream of the β3-AR-cAMP-PKA-p38 (51,52). To investigate this, we performed an unbiased DiffTF analysis (65) of our RNA-seq data to identified the TFs responsible for the gene expression patterns observed in Fig. 1. This analysis (see Methods) predicted that SREBP1, PPARA, and PPARG were the primary active TFs 22 days after SUMOylation inhibition (Fig. 6A). Consistently, beiging genes such as *SLC2A4* (GLUT4), *LEP* (Leptin) and *UCP1* were predicted targets of PPARA and PPARG, *ACSL1* (Acyl-CoA Synthetase) of PPARA, and *ADIPOR2* (Adiponectin receptor) of PPARG upon transient SUMOylation inhibition (Fig. 6B and C). These findings suggest that upregulation of CEBPs and the cAMP-PKA-p38 pathway led to PPARA/G activation. Alternatively, SUMOylation inhibition may directly impact PPARA and PPARG transactivation activity, given that both are well-characterized SUMO targets, and SUMOylation can either enhance or repress their function (5,15,98,99). Based on this, and since *UCP1* is not a direct target of SREBP1 (Suppl. Fig. 6A), we focused our attention on PPARA and PPARG activation.

As beiging features reprogramming of PPARG enhancers in human cells (76), we sought to validate the activation of PPARG in response to TAK-981 by measuring the activity of PPAR response elements (PPREs) linked to key beiging genes: *TSHR*, *ACSL1*, *UCP1*, *PPARGC1A*, and *CIDEA* (Suppl. Fig. 6B-F). These PPREs were primarily identified using the JASPAR database (75). To assess PPRE activity, we performed ChIP-qPCR for the occurrence of H3K27ac (active enhancers) and H3K27me3 (repressed enhancers) at day 22 post adipogenic induction (Fig. 1A). These experiments showed that treatment with either rosiglitazone or TAK-981 alone did not strongly increase the occurrence of H3K27ac at *TSHR* and *ACSL1* PPREs (Fig. 6D, left). At the same time, we observed a decrease in H3K27me3, which was particularly evident at *^PPRE1^TSHR* and *^PPRE^ACSL1* (Fig. 6E, left). Importantly, when rosiglitazone and TAK-981 were combined, the increase in H3K27ac levels became more significant at these PPREs, while the presence of H3K27me3 also decreased significantly (Fig. 6D and E, left). These support a previous observation that *TSHR* expression is directly linked to *UCP1* activation during adaptive thermogenesis (88).

When examining *UCP1* PPREs (Suppl. Fig. 6F), we found that TAK-981 treatment alone resulted in a strong and significant increase in H3K27ac, accompanied by a decrease in H3K27me3 while rosiglitazone did not affect these marks (Fig. 6F,G. However, co-treatment with rosiglitazone and TAK-981 significantly increased further H3K27ac occurrence while H3K27me3 remained low (Fig. 6F,G). Importantly, the levels of H3K27ac gradually increased from *^PPRE1^UCP1* to *^PPRE5^UCP1*, a pattern consistent with previous PPARG and H3K27ac ChIP-seq data from beige adipocytes (Suppl. Fig. 6G) (76) and reflecting our *UCP1* expression data. Similar trends were observed when assessing *PPARGC1A* and *CIDEA* PPREs, although the effects of TAK-981 and rosiglitazone were less pronounced than for *UCP1 or ^PPRE1^TSHR*, particularly for H3K27ac at the PPRE of *CIDEA* (Fig. 6D,E).

We next hypothesized that increased *PPARG* mRNA levels (Fig. 2A,C) and protein levels (Fig. 5I) mediated the positive effect of SUMOylation inhibition on beiging genes transactivation, including *UCP1*. We thus measured PPARG occurrence at *UCP1* PPREs by ChIP-qPCR 21 days after adipogenic induction. We could not reliably detect PPARG at *^PPRE1^UCP1,* which may be expected from previous PPARG ChIP-seq experiments (Suppl. Fig. 6G) (76). However, we observed a significant increase of PPARG detection at *^PPRE2^UCP1* to *^PPRE4^UCP1* and an increasing trend at *^PPRE5^UCP1* in cells co-treated with TAK-981 and rosiglitazone while other treatments did not have notable effects (Fig. 6G).

Together, these data are consistent with a model in which transient inhibition of SUMOylation primed and stably facilitated the epigenetic activation of PPREs at thermogenic enhancers and stably enhanced PPARG protein levels to promote adaptive thermogenesis.

## Discussion

The cellular mechanisms that govern the phenotypic plasticity and fate determination between white to beige adipocytes remain poorly understood. In this study we revealed a key role for SUMOylation in this process. However, while we have recently shown that low SUMOylation levels drive towards adipogenic rather than osteogenic transcription in mouse beige adipocytes (81), inhibiting SUMOylation was not sufficient to cause beiging in human pre-adipocytes, unless combined with the PPARG agonist rosiglitazone. This synergy between TAK-981 and rosiglitazone, which we observed both in a pre-adipocyte cell line and primary hASCs, was evidenced by substantial enhancer and transcriptional reprogramming driven by CEBPs and PPARG, activation of the β3-adrenergique pathway, increased expression of the thermogenic protein UCP1 and subsequent mitochondrial uncoupling. These results are consistent with a model in which SUMOylation serves as a barrier against transcriptional reprogramming to maintain white cell’s identity.

WAT plasticity allows adaptive thermogenesis, which may occur via two routes: either through *de novo* differentiation of white/beige precursors into beige cells or the reversible conversion of white cells into beige cells (27,28,100,101). Whether one of these routes is favored during hypothermia or whether both contribute to beiging remains a topic of debate. In this study, transient SUMOylation inhibition in white pre-adipocytes enabled *de novo* beiging in the presence of rosiglitazone. This strongly suggests that beiging transcription programs are silent yet epigenetically imprinted in pre-adipocytes and retained in mature adipocytes. The transient TAK-981 treatment may have created an epigenetic priming in pre-adipocytes and hASCs, allowing them to response to rosiglitazone and trigger expression of beiging genes like UCP1. When SUMOylation recovers, it fulfils its function of guardian of cellular identity by maintaining the newly acquired beige identity (Fig. 6H). Future work should now investigate whether SUMOylation inhibition in mature adipocytes can induce the conversion towards a beige phenotype as efficiently as de novo beige differentiation.

Beiging has been shown to be reversible, i.e. beige cells can revert to white adipocytes upon warming (101). However, we found that inhibition of SUMOylation by TAK-981 imprinted a stable beiging fate that was not reversed by washing out the drug or by introducing a pause in the rosiglitazone treatment. This observation suggests that temporal relief of the SUMO barrier allows cells to assume a new equilibrium that may involve implementation of a stable beige adipocyte epigenetic memory. This is consistent with our recent findings that SUMOylation also preserves and prevents osteoblast fate in adipocytes, further underscoring the role of SUMOylation in regulating adipose tissue identity, plasticity, and function (81).

Why does SUMOylation inhibition specifically promote beiging in the presence of rosiglitazone? Both our data and prior reports indicate that blocking SUMOylation commonly elicits a remarkably monomorphic, cell-type-specific transcriptional outcome (102). Recent mechanistic work at the *IFNB1* locus provides a clear example of how this can occur: SUMOylation inhibition alone is sufficient to activate *IFNB1* expression, accompanied by coordinated changes in chromatin accessibility, histone modifications, and three-dimensional enhancer-promoter contacts, even in the absence of infection (91). This locus-centric mechanism implies that SUMOylation acts upstream of highly constrained regulatory programs centered on individual, functionally dominant genes. By analogy, the beiging phenotype may arise because SUMOylation inhibition, together with rosiglitazone, selectively trigger a cascade of local chromatin remodeling and enhancer activation to derepress the *UCP1* regulatory unit, which shifts adipocyte metabolism from energy storage to energy dissipation. Thus, focused dissection of single, dominant loci such as *IFNB1* or *UCP1* is a powerful strategy to understand how SUMOylation controls apparently monomorphic, cell-type specific responses.

Although the transcriptomes of white and beige adipocytes largely overlap, and UCP1 uncoupling activity is a hallmark of beiging, adaptive thermogenesis still involves extensive transcriptional remodeling to support the metabolic shift from energy storage to energy expenditure. This process is driven by the activation of beige-specific genes such as *UCP1*, *CIDEA* and *ELOVL3* (92). Using the BATLAS as a reference, we show that transient SUMOylation inhibition stably shifts the transcriptome toward beiging in a rosiglitazone-dependent manner. The ability of transient hypoSUMOylation to induce a stable transcriptional and cellular fate change suggests that SUMOylation plays a key role in epigenetic regulation in pre-adipocytes and mature adipocytes, as previously shown in immune cells, ESCs and beige cells (2,81,91). Specifically, transient SUMOylation inhibition increased the accessibility to several enhancers, including those bound by early adipogenic factors like CEBPs (Fig. 6H). CEBPB is particularly mobilized early on, which is supported by increased mRNA and protein levels, stabilized by deSUMOylation, and promotes the expression of downstream targets CEBPA and PPARG (Fig. 6G) (103). This is evidenced by increased expression of *CEBPA* and *PPARG*, especially the beige-specific isoform PPARG2, as well as strong accessibility of their cognate enhancers toward the end of the time course. As shown previously, these factors then promote each-other’s expression via a positive feedback loop and support the activation of the beiging transcriptional program (76,103). This implies that SUMOylation actively enforces beiging gene silencing at the level of enhancers in pre-adipocytes to limit adaptive thermogenesis and response to PPARG agonists. In support, our study of thermogenic enhancers suggests that while SUMOylation inhibition primarily affects chromatin accessibility by facilitating histone acetylation at *UCP1* PPREs, the presence of rosiglitazone is necessary to achieve maximal *UCP1* expression. This supports a model in which SUMOylation inhibition leads to epigenetic remodeling, whereas rosiglitazone enhances *UCP1* transactivation via PPARG (Fig. 6G).

How might SUMOylation work as a barrier in white pre-adipocytes and primary hASCs at the molecular level? Prior mass spectrometry experiments in adipocytes and other cell types have suggested that SUMOylation of chromatin-bound proteins within heterochromatin-maintaining complexes restricts adaptive gene activation and transcriptional reprogramming (5,83,96,104,105). Furthermore, our data reveal that transient SUMOylation inhibition in pre-adipocytes leads to a reduction in H3K27me3 at beiging enhancers even after SUMOylation recovery, suggesting that TAK-981 disrupts Polycomb-mediated histone methylation, ultimately establishing a stable epigenetic beiging fate. A similar link between PRC2, H3K27me3, SUMOylation, and energy metabolism preservation in adipocytes and cancer cells has been proposed (106,107). These findings mirror a recent study in *Drosophila*, where transient Polycomb downregulation resulted in stable epigenetic reprogramming toward a cancerous state (108). This raises the possibility that SUMOylation serves as a key regulator of Polycomb, maintaining H3K27me3 homeostasis to prevent epigenetic drift towards unwanted identity transitions or into disease states. These insights are also relevant to energy storage in obesity, which has been recently linked to an epigenetic obesogenic memory (35).

In parallel, we observed an increase in H3K27ac levels at beiging enhancers depleted of H3K27me3, suggesting that SUMOylation inhibition facilitates histone acetylation, a process known to contribute to *UCP1* expression via HAT1 (109). Consistently, recent studies have identified HATs as major SUMO targets across various cell types, reinforcing the idea that SUMOylation regulates histone acetylation (5,83,96,98,104). SUMOylation can inhibit p300, often acting as a transcriptional repressor, by blocking its ability to acetylate histones. This occurs through mechanisms like the SUMOylation of p300 itself or by modifying other proteins like the H4 histone tail, which then physically interferes with p300’s activity (110,111). The presence of SUMO modifications on p300 or its targets can shift the balance from transcriptional activation to repression. Further investigation using techniques such as ChIP-sequencing, RIME (83) or PLAM-seq (105) is needed to determine the nature of SUMOylated epigenetic regulators responsible for repressing beiging fate in pre-adipocytes. This is the focus of a follow up study.

Increased expression of UCP1 upon treatment with rosiglitazone and TAK-981 was reflected in a metabolic shift with higher OCR and ECAR. As reported by others, UCP1-induced OCR was not correlated with increased mitochondrial ATP production, but instead increased thermogenesis (112). Rather, we observed increased glycolysis associated with increased ATP production. This phenotype suggests that differentiated cells display a glycolytic and high energy-demanding phenotype, as expected from beige cells (76,113). Interestingly, cells differentiated in the presence of rosiglitazone and TAK-981 were almost as glycolytic as brown adipocytes. This suggests that rosiglitazone together with TAK-981 induces a highly energy producing fat cell phenotype which may prompt to explore their combination for therapeutic use *in vivo*.

During adaptive thermogenesis, activation of the β3-AR-cAMP-PKA-p38 signaling pathway mobilizes specific transcription factors, including PPARA/G and coactivators to induce the expression of beiging genes like *UCP1*. Our data suggest that SUMOylation inhibition activates this pathway at two levels. RNA sequencing revealed that SUMOylation facilitates the repression of genes involved in this signaling cascade, though the precise mechanism remains unclear. Additionally, western blot analysis of the phosphorylation of PKA and p38 substrates suggests that SUMOylation may also suppress the activity of these kinases. Since both PKA (catalytic and regulatory subunits) and p38 have been identified as SUMOylation targets (5,96,98,114,115) (Suppl. Table 6), SUMOylation could directly inhibit their enzymatic activity or modify their substrate specificity. Moreover, although the β3-AR has not been identified as a SUMO target yet, many other proteins of the thermogenic pathway, including adenylate cyclase, OXPHOS proteins, PKA and p38 targets and lipolytic enzymes have be shown to be SUMO targets across different tissues and species using mass spectrometry (5,96,98,114,115) (Suppl. Table 6). These findings support a model in which SUMOylation acts as a regulatory constraint on cAMP-PKA-p38 signaling at both the transcription and protein levels, thereby restricting the initiation of beiging. However, the relative contribution of the SUMOylation pathway to cAMP signaling versus epigenetics in beiging remains to be evaluated.

SUMOylation may also directly modulate transcriptional regulators. It has been shown to restrict the transactivation activity of TFs like CEBPB and PPARG (12,81,99,113). We found that CEBPB SUMOylation strongly decreases upon TAK-981 treatment, which has been shown to boost its transactivation activity (12). In addition, TAK-981 may directly enhance the transcriptional activity of PPARG, increasing the expression of key beiging genes like *UCP1* (99). Furthermore, recent studies suggest that preventing the SUMOylation of transcriptional activators, including nuclear receptors and coactivators, can alter their chromatin localization (5,14,83,104,116,117). Our recent findings show that chronic SUMOylation inhibition in differentiating 3T3-L1 cells disrupts the chromatin occupancy of RXR, leading to significant changes in transcriptional output (5). Given that rosiglitazone is a PPARG agonist, its effect on chromatin localization, along with other activators and coactivators, may be influenced by TAK-981. This, and other studies of the glucocorticoid receptor (83,104), suggests that TAK-981 could help reprogram the transcriptome toward beiging by stabilizing the recruitment of PPAR-RXR and coactivators like epigenetic factors at enhancers like those controlling *UCP1*.

It is worth mentioning that although our findings are in accordance with previous links drawn between genetic manipulation of SUMOylation enzymes and energy homeostasis in adipocytes (118) and the fact that boosting SUMOylation drives osteogenesis rather than adipogenesis in adipose precursors (81), they are also in apparent contrast with studies in WAT-specific SENP2 knockout (KO) mice (119). These mice lack the deSUMOylase SENP2, leading to increased SUMOylation levels of certain proteins, such as CEBPB. The increased SUMOylation of CEBPB inhibits its function, resulting in lower expression of HOXC10, a browning inhibitor, through recruitment of the transcriptional repressor DAXX. Therefore, higher SUMOylation levels in WAT enhanced beiging capacity (119). We do not presently understand why we observed the opposite phenotype, where loss of SUMOylation leads to increased beiging. One possible explanation could be differences in the dynamics of the process. In cell lacking SENP2, SUMO levels are chronically increased in both pre-adipocytes and mature adipocytes, which may have a much wider impact on transcriptional programs, including secondary effects and activation of compensatory mechanisms that may not be operative under physiological conditions. It is also possible that SUMO serves as a rheostat, in which either too little SUMOylation or too much SUMOylation lead to the same phenotype. Clearly, more work is needed to clarify these differences. Nonetheless, from a therapeutic perspective our approach using TAK-981 offers a distinct advantage. As a small-molecule inhibitor, TAK-981 can be used transiently to induce reprogramming of adipose tissue, which may be tested *in vivo*.

In this study, we characterized the stable transcriptional and cellular identity shifts initiated by transient SUMOylation inhibition, as well as alterations of chromatin modifications and accessibility at selected beiging enhancers several days after TAK-981 treatment. Our findings suggest that TAK-981 imprints a stable beiging fate, priming cells to respond more robustly to rosiglitazone. Given that derepression of *PPARG* and *UCP1* also occurred in primary hASCs, these findings may provide new inroads in treatment of metabolic disorders using carefully defined sequential drug treatment algorithms. Furthermore, our data broaden the involvement of SUMOylation in preventing epigenetic drift, potentially safeguarding against disease.

## Acknowledgements

We thank Dr. Damien Dufour, Prof. Simon N. Dankel, Prof. Gunnar Mellgren, Dr. Jan-Inge Bjune, Dr. Sandra Lopez-Aviles for their input. Sequencing was performed by the GenomEast platform, a member of the ‘France Génomique’ consortium (ANR-10-INBS-0009). We thank Dr. Stéphanie Le Gras for supporting RNA-seq data analysis.

## Data availability

RNA-and ATAC-sequencing data are available at the Gene Expression Omnibus using accession number GSE290926 and GSE313330, respectively. H3K27ac and PPARG ChIP-seq datasets were downloaded from the GEO using accession number GSM1443810, GSM1443812, GSM1443806 and GSM1443808.

## Author contributions

**Patrizia Maria Christiane Nothnagel:** Investigation, Formal analysis, Visualization, Writing - Original Draft. **Paul-Arthur Meslin:** Investigation, Investigation, Formal analysis, Visualization. **Jonas Aakre Wik:** Methodology, Investigation. **Yunna E. Strøm:** Investigation. **Magnar Bjørås:** Funding acquisition, Resources. **Jorrit M. Enserink:** Resources, Writing - Review & Editing. **Bjørn Steen Skålhegg:** Supervision, Resources, Methodology, Writing - Review & Editing. **Nolwenn Briand:** Supervision, Resources, Methodology, Investigation. **Anthony Mathelier:** Supervision, Methodology, Investigation, Formal analysis, Visualization, Funding acquisition, Writing - Review & Editing. **Pierre Chymkowitch:** Conceptualization, Methodology, Resources, Supervision, Funding acquisition, Writing - Original Draft, Writing - Review & Editing.

## Footnotes

1 Global Burden of Disease Collaborative Network. Global Burden of Disease Study 2021 (GBD 2021). Seattle, United States: Institute for Health Metrics and Evaluation (IHME), 2024.

## References

1. Chymkowitch, P., Nguea, P.A. and Enserink, J.M. (2015) SUMO-regulated transcription: challenging the dogma. Bioessays, 37, 1095–1105.

2. Cossec, J.C., Theurillat, I., Chica, C., Bua Aguin, S., Gaume, X., Andrieux, A., Iturbide, A., Jouvion, G., Li, H., Bossis, G. et al. (2018) SUMO Safeguards Somatic and Pluripotent Cell Identities by Enforcing Distinct Chromatin States. Cell Stem Cell, 23, 742–757 e748.

3. Cossec, J.C., Traboulsi, T., Sart, S., Loe-Mie, Y., Guthmann, M., Hendriks, I.A., Theurillat, I., Nielsen, M.L., Torres-Padilla, M.E., Baroud, C.N. et al. (2023) Transient suppression of SUMOylation in embryonic stem cells generates embryo-like structures. Cell reports, 42, 112380.

4. Dufour, D., Dumontet, T., Sahut-Barnola, I., Carusi, A., Onzon, M., Pussard, E., Wilmouth, J.J., Olabe, J., Lucas, C., Levasseur, A. et al. (2022) Loss of SUMO-specific protease 2 causes isolated glucocorticoid deficiency by blocking adrenal cortex zonal transdifferentiation in mice. Nat Commun, 13, 7858.

5. Zhao, X., Hendriks, I.A., Le Gras, S., Ye, T., Ramos-Alonso, L., Nguea, P.A., Lien, G.F., Ghasemi, F., Klungland, A., Jost, B., et al. (2022) Waves of sumoylation support transcription dynamics during adipocyte differentiation. Nucleic Acids Res, 50, 1351–1369.

6. Pei, J., Zou, D., Li, L., Kang, L., Sun, M., Li, X., Chen, Q., Chen, D., Qu, B., Gao, X. et al. (2024) Senp7 deficiency impairs lipid droplets maturation in white adipose tissues via Plin4 deSUMOylation. Journal of Biological Chemistry, 300, 107319.

7. Demarque, M.D., Nacerddine, K., Neyret-Kahn, H., Andrieux, A., Danenberg, E., Jouvion, G., Bomme, P., Hamard, G., Romagnolo, B., Terris, B. et al. (2011) Sumoylation by Ubc9 regulates the stem cell compartment and structure and function of the intestinal epithelium in mice. Gastroenterology, 140, 286–296.

8. Mikkonen, L., Hirvonen, J. and Janne, O.A. (2013) SUMO-1 regulates body weight and adipogenesis via PPARgamma in male and female mice. Endocrinology, 154, 698–708.

9. Zheng, Q., Cao, Y., Chen, Y., Wang, J., Fan, Q., Huang, X., Wang, Y., Wang, T., Wang, X., Ma, J. et al. (2018) Senp2 regulates adipose lipid storage by de-SUMOylation of Setdb1. J Mol Cell Biol, 10, 258–266.

10. Cignarelli, A., Melchiorre, M., Peschechera, A., Conserva, A., Renna, L.A., Miccoli, S., Natalicchio, A., Perrini, S., Laviola, L. and Giorgino, F. (2010) Role of UBC9 in the regulation of the adipogenic program in 3T3-L1 adipocytes. Endocrinology, 151, 5255–5266.

11. Liu, B., Wang, T., Mei, W., Li, D., Cai, R., Zuo, Y. and Cheng, J. (2014) Small ubiquitin-like modifier (SUMO) protein-specific protease 1 de-SUMOylates Sharp-1 protein and controls adipocyte differentiation. J Biol Chem, 289, 22358–22364.

12. Chung, S.S., Ahn, B.Y., Kim, M., Choi, H.H., Park, H.S., Kang, S., Park, S.G., Kim, Y.-B., Cho, Y.M., Lee, H.K. et al. (2010) Control of Adipogenesis by the SUMO-Specific Protease SENP2. Molecular and Cellular Biology, 30, 2135–2146.

13. Antonio Urrutia, G., Ramachandran, H., Cauchy, P., Boo, K., Ramamoorthy, S., Boller, S., Dogan, E., Clapes, T., Trompouki, E., Torres-Padilla, M.-E., et al. (2021) ZFP451-mediated SUMOylation of SATB2 drives embryonic stem cell differentiation. Genes & Development, 35, 1142–1160.

14. Rosonina, E., Akhter, A., Dou, Y., Babu, J. and Sri Theivakadadcham, V.S. (2017) Regulation of transcription factors by sumoylation. Transcription, 8, 220–231.

15. Boulanger, M., Chakraborty, M., Tempe, D., Piechaczyk, M. and Bossis, G. (2021) SUMO and Transcriptional Regulation: The Lessons of Large-Scale Proteomic, Modifomic and Genomic Studies. Molecules, 26.

16. Seeler, J.S. and Dejean, A. (2017) SUMO and the robustness of cancer. Nat Rev Cancer, 17, 184–197.

17. Dufour, D., Zhao, X., Chaleil, F., Nothnagel, P.M.C., Bjørås, M., Lefrançois-Martinez, A.-M., Martinez, A. and Chymkowitch, P. (2025) Pharmacological inhibition of SUMOylation with TAK-981 mimics genetic HypoSUMOylation in murine perigonadal white adipose tissue. Adipocyte, 14, 2474107.

18. Sapir, A. (2020) Not So Slim Anymore-Evidence for the Role of SUMO in the Regulation of Lipid Metabolism. Biomolecules, 10.

19. Shapira, S.N. and Seale, P. (2019) Transcriptional Control of Brown and Beige Fat Development and Function. Obesity (Silver Spring), 27, 13–21.

20. Rosenwald, M. and Wolfrum, C. (2014) The origin and definition of brite versus white and classical brown adipocytes. Adipocyte, 3, 4–9.

21. Chang, S.H., Song, N.J., Choi, J.H., Yun, U.J. and Park, K.W. (2019) Mechanisms underlying UCP1 dependent and independent adipocyte thermogenesis. Obes Rev, 20, 241–251.

22. Christy, R.J., Kaestner, K.H., Geiman, D.E. and Lane, M.D. (1991) CCAAT/enhancer binding protein gene promoter: binding of nuclear factors during differentiation of 3T3-L1 preadipocytes. Proc Natl Acad Sci U S A, 88, 2593–2597.

23. Clarke, S.L., Robinson, C.E. and Gimble, J.M. (1997) CAAT/enhancer binding proteins directly modulate transcription from the peroxisome proliferator-activated receptor gamma 2 promoter. Biochem Biophys Res Commun, 240, 99–103.

24. Lee, C.H., Olson, P. and Evans, R.M. (2003) Minireview: lipid metabolism, metabolic diseases, and peroxisome proliferator-activated receptors. Endocrinology, 144, 2201–2207.

25. Farmer, S.R. (2006) Transcriptional control of adipocyte formation. Cell Metab, 4, 263–273.

26. Rosenwald, M., Perdikari, A., Rulicke, T. and Wolfrum, C. (2013) Bi-directional interconversion of brite and white adipocytes. Nat Cell Biol, 15, 659–667.

27. Merrick, D., Sakers, A., Irgebay, Z., Okada, C., Calvert, C., Morley, M.P., Percec, I. and Seale, P. (2019) Identification of a mesenchymal progenitor cell hierarchy in adipose tissue. Science, 364.

28. Schwalie, P.C., Dong, H., Zachara, M., Russeil, J., Alpern, D., Akchiche, N., Caprara, C., Sun, W., Schlaudraff, K.U., Soldati, G. et al. (2018) A stromal cell population that inhibits adipogenesis in mammalian fat depots. Nature, 559, 103–108.

29. Pahlavani, M., Pham, K., Kalupahana, N.S., Morovati, A., Ramalingam, L., Abidi, H., Kiridana, V. and Moustaid-Moussa, N. (2024) Thermogenic adipose tissues: Promising therapeutic targets for metabolic diseases. J Nutr Biochem, 137, 109832.

30. Liu, X.Z., Pedersen, L. and Halberg, N. (2021) Cellular mechanisms linking cancers to obesity. Cell Stress, 5, 55–72.

31. Pati, S., Irfan, W., Jameel, A., Ahmed, S. and Shahid, R.K. (2023) Obesity and Cancer: A Current Overview of Epidemiology, Pathogenesis, Outcomes, and Management. Cancers (Basel), 15.

32. Nguyen, H.L., Geukens, T., Maetens, M., Aparicio, S., Bassez, A., Borg, A., Brock, J., Broeks, A., Caldas, C., Cardoso, F. et al. (2023) Obesity-associated changes in molecular biology of primary breast cancer. Nat Commun, 14, 4418.

33. Nordsletten, M., Saeed, U., Myklebust, T., Robsahm, T.E., Møller, B., Skålhegg, B.S., Mala, T. and Yaqub, S. (2023) Body mass index and its association with 22 cancer types: a Norwegian cohort study of 481 202 cancer cases. Acta Oncol, 62, 1273–1278.

34. Kalligeros, M., Shehadeh, F., Mylona, E.K., Benitez, G., Beckwith, C.G., Chan, P.A. and Mylonakis, E. (2020) Association of Obesity with Disease Severity among Patients with COVID-19. Obesity (Silver Spring).

35. Hinte, L.C., Castellano-Castillo, D., Ghosh, A., Melrose, K., Gasser, E., Noe, F., Massier, L., Dong, H., Sun, W., Hoffmann, A. et al. (2024) Adipose tissue retains an epigenetic memory of obesity after weight loss. Nature.

36. Hanssen, M.J., Hoeks, J., Brans, B., van der Lans, A.A., Schaart, G., van den Driessche, J.J., Jorgensen, J.A., Boekschoten, M.V., Hesselink, M.K., Havekes, B., et al. (2015) Short-term cold acclimation improves insulin sensitivity in patients with type 2 diabetes mellitus. Nat Med, 21, 863–865.

37. Herbers, E., Patrikoski, M., Wagner, A., Jokinen, R., Hassinen, A., Heinonen, S., Miettinen, S., Peltoniemi, H., Pirinen, E. and Pietilainen, K.H. (2022) Preventing White Adipocyte Browning during Differentiation In Vitro: The Effect of Differentiation Protocols on Metabolic and Mitochondrial Phenotypes. Stem Cells Int, 2022, 3308194.

38. Wang, H.J., Lee, C.S., Yee, R.S.Z., Groom, L., Friedman, I., Babcock, L., Georgiou, D.K., Hong, J., Hanna, A.D., Recio, J. et al. (2020) Adaptive thermogenesis enhances the life-threatening response to heat in mice with an Ryr1 mutation. Nat Commun, 11, 5099.

39. Potolitsyna, E., Pickering, S.H., Bellanger, A., Germier, T., Collas, P. and Briand, N. (2024) Cytoskeletal rearrangement precedes nucleolar remodeling during adipogenesis. Commun Biol, 7, 458.

40. Hazell Pickering, S., Abdelhalim, M., Collas, P. and Briand, N. (2024) Alternative isoform expression of key thermogenic genes in human beige adipocytes. Front Endocrinol (Lausanne), 15, 1395750.

41. Hazell Pickering, S., Galigniana, N.M., Abdelhalim, M., Sørensen, A.L., Zucknick, M., Collas, P. and Briand, N. (2025) Multimodal epigenetic and enhancer network remodeling shape the transcriptional landscape of beige adipocytes. bioRxiv, 2025.2003.2028.645896.

42. Hazell Pickering, S., Abdelhalim, M., Collas, P. and Briand, N. (2024) Alternative isoform expression of key thermogenic genes in human beige adipocytes. Frontiers in Endocrinology, 15.

43. Martin, M. (2011) Cutadapt removes adapter sequences from high-throughput sequencing reads. 2011, 17, 3.

44. Langmead, B. and Salzberg, S.L. (2012) Fast gapped-read alignment with Bowtie 2. Nature methods, 9, 357–359.

45. Dobin, A., Davis, C.A., Schlesinger, F., Drenkow, J., Zaleski, C., Jha, S., Batut, P., Chaisson, M. and Gingeras, T.R. (2013) STAR: ultrafast universal RNA-seq aligner. Bioinformatics, 29, 15–21.

46. Anders, S., Pyl, P.T. and Huber, W. (2015) HTSeq--a Python framework to work with high-throughput sequencing data. Bioinformatics, 31, 166–169.

47. Love, M.I., Huber, W. and Anders, S. (2014) Moderated estimation of fold change and dispersion for RNA-seq data with DESeq2. Genome biology, 15, 550.

48. Perdikari, A., Leparc, G.G., Balaz, M., Pires, N.D., Lidell, M.E., Sun, W., Fernandez-Albert, F., Müller, S., Akchiche, N., Dong, H. et al. (2018) BATLAS: Deconvoluting Brown Adipose Tissue. Cell reports, 25, 784–797.e784.

49. Durinck, S., Spellman, P.T., Birney, E. and Huber, W. (2009) Mapping identifiers for the integration of genomic datasets with the R/Bioconductor package biomaRt. Nature protocols, 4, 1184–1191.

50. Yu, G., Wang, L.G., Han, Y. and He, Q.Y. (2012) clusterProfiler: an R package for comparing biological themes among gene clusters. OMICS, 16, 284–287.

51. Badia, I.M.P., Velez Santiago, J., Braunger, J., Geiss, C., Dimitrov, D., Muller-Dott, S., Taus, P., Dugourd, A., Holland, C.H., Ramirez Flores, R.O., et al. (2022) decoupleR: ensemble of computational methods to infer biological activities from omics data. Bioinform Adv, 2, vbac016.

52. Turei, D., Valdeolivas, A., Gul, L., Palacio-Escat, N., Klein, M., Ivanova, O., Olbei, M., Gabor, A., Theis, F., Modos, D. et al. (2021) Integrated intra-and intercellular signaling knowledge for multicellular omics analysis. Mol Syst Biol, 17, e9923.

53. Buenrostro, J.D., Wu, B., Chang, H.Y. and Greenleaf, W.J. (2015) ATAC-seq: A Method for Assaying Chromatin Accessibility Genome-Wide. Current Protocols in Molecular Biology, 109.

54. Corces, M.R., Trevino, A.E., Hamilton, E.G., Greenside, P.G., Sinnott-Armstrong, N.A., Vesuna, S., Satpathy, A.T., Rubin, A.J., Montine, K.S., Wu, B. et al. (2017) An improved ATAC-seq protocol reduces background and enables interrogation of frozen tissues. Nature methods, 14, 959–962.

55. Ramos-Alonso, L., Holland, P., Le Gras, S., Zhao, X., Jost, B., Bjørås, M., Barral, Y., Enserink, J.M. and Chymkowitch, P. (2023) Mitotic chromosome condensation resets chromatin to safeguard transcriptional homeostasis during interphase. Proceedings of the National Academy of Sciences, 120.

56. Buenrostro, J.D., Giresi, P.G., Zaba, L.C., Chang, H.Y. and Greenleaf, W.J. (2013) Transposition of native chromatin for fast and sensitive epigenomic profiling of open chromatin, DNA-binding proteins and nucleosome position. Nature methods, 10, 1213–1218.

57. Li, H., Handsaker, B., Wysoker, A., Fennell, T., Ruan, J., Homer, N., Marth, G., Abecasis, G., Durbin, R. and Genome Project Data Processing, S. (2009) The Sequence Alignment/Map format and SAMtools. Bioinformatics, 25, 2078–2079.

58. Ramirez, F., Ryan, D.P., Gruning, B., Bhardwaj, V., Kilpert, F., Richter, A.S., Heyne, S., Dundar, F. and Manke, T. (2016) deepTools2: a next generation web server for deep-sequencing data analysis. Nucleic Acids Res, 44, W160–165.

59. Zhang, Y., Liu, T., Meyer, C.A., Eeckhoute, J., Johnson, D.S., Bernstein, B.E., Nusbaum, C., Myers, R.M., Brown, M., Li, W. et al. (2008) Model-based analysis of ChIP-Seq (MACS). Genome biology, 9, R137.

60. Heinz, S., Benner, C., Spann, N., Bertolino, E., Lin, Y.C., Laslo, P., Cheng, J.X., Murre, C., Singh, H. and Glass, C.K. (2010) Simple combinations of lineage-determining transcription factors prime cis-regulatory elements required for macrophage and B cell identities. Molecular cell, 38, 576–589.

61. Quinlan, A.R. and Hall, I.M. (2010) BEDTools: a flexible suite of utilities for comparing genomic features. Bioinformatics, 26, 841–842.

62. Anders, S. and Huber, W. (2010) Differential expression analysis for sequence count data. Genome biology, 11, R106.

63. Kumar, L. and M, E.F. (2007) Mfuzz: a software package for soft clustering of microarray data. Bioinformation, 2, 5–7.

64. Welch, R.P., Lee, C., Imbriano, P.M., Patil, S., Weymouth, T.E., Smith, R.A., Scott, L.J. and Sartor, M.A. (2014) ChIP-Enrich: gene set enrichment testing for ChIP-seq data. Nucleic Acids Research, 42, e105–e105.

65. Berest, I., Arnold, C., Reyes-Palomares, A., Palla, G., Rasmussen, K.D., Giles, H., Bruch, P.-M., Huber, W., Dietrich, S., Helin, K. et al. (2019) Quantification of Differential Transcription Factor Activity and Multiomics-Based Classification into Activators and Repressors: diffTF. Cell reports, 29, 3147–3159.e3112.

66. Puig, R.R., Boddie, P., Khan, A., Castro-Mondragon, J.A. and Mathelier, A. (2021) UniBind: maps of high-confidence direct TF-DNA interactions across nine species. BMC genomics, 22, 482.

67. Luo, W. and Brouwer, C. (2013) Pathview: an R/Bioconductor package for pathway-based data integration and visualization. Bioinformatics, 29, 1830–1831.

68. Schindelin, J., Arganda-Carreras, I., Frise, E., Kaynig, V., Longair, M., Pietzsch, T., Preibisch, S., Rueden, C., Saalfeld, S., Schmid, B. et al. (2012) Fiji: an open-source platform for biological-image analysis. Nature methods, 9, 676–682.

69. Cantelmo, A.R., Conradi, L.C., Brajic, A., Goveia, J., Kalucka, J., Pircher, A., Chaturvedi, P., Hol, J., Thienpont, B., Teuwen, L.A. et al. (2016) Inhibition of the Glycolytic Activator PFKFB3 in Endothelium Induces Tumor Vessel Normalization, Impairs Metastasis, and Improves Chemotherapy. Cancer Cell, 30, 968–985.

70. Corces, M.R., Trevino, A.E., Hamilton, E.G., Greenside, P.G., Sinnott-Armstrong, N.A., Vesuna, S., Satpathy, A.T., Rubin, A.J., Montine, K.S., Wu, B. et al. (2017) An improved ATAC-seq protocol reduces background and enables interrogation of frozen tissues. Nature methods, 14, 959–962.

71. Kent, W.J., Sugnet, C.W., Furey, T.S., Roskin, K.M., Pringle, T.H., Zahler, A.M. and Haussler, D. (2002) The human genome browser at UCSC. Genome Res, 12, 996–1006.

72. Frankish, A., Carbonell-Sala, S., Diekhans, M., Jungreis, I., Loveland, J.E., Mudge, J.M., Sisu, C., Wright, J.C., Arnan, C., Barnes, I. et al. (2023) GENCODE: reference annotation for the human and mouse genomes in 2023. Nucleic Acids Res, 51, D942–D949.

73. Pruitt, K.D., Brown, G.R., Hiatt, S.M., Thibaud-Nissen, F., Astashyn, A., Ermolaeva, O., Farrell, C.M., Hart, J., Landrum, M.J., McGarvey, K.M. et al. (2014) RefSeq: an update on mammalian reference sequences. Nucleic Acids Res, 42, D756–763.

74. Tweedie, S., Braschi, B., Gray, K., Jones, T.E.M., Seal, R.L., Yates, B. and Bruford, E.A. (2021) Genenames.org: the HGNC and VGNC resources in 2021. Nucleic Acids Res, 49, D939–D946.

75. Rauluseviciute, I., Riudavets-Puig, R., Blanc-Mathieu, R., Castro-Mondragon, J.A., Ferenc, K., Kumar, V., Lemma, R.B., Lucas, J., Chèneby, J., Baranasic, D. et al. (2024) JASPAR 2024: 20th anniversary of the open-access database of transcription factor binding profiles. Nucleic Acids Res, 52, D174–d182.

76. Loft, A., Forss, I., Siersbæk, M.S., Schmidt, S.F., Larsen, A.S., Madsen, J.G., Pisani, D.F., Nielsen, R., Aagaard, M.M., Mathison, A. et al. (2015) Browning of human adipocytes requires KLF11 and reprogramming of PPARγ superenhancers. Genes Dev, 29, 7–22.

77. Xue, R., Lynes, M.D., Dreyfuss, J.M., Shamsi, F., Schulz, T.J., Zhang, H., Huang, T.L., Townsend, K.L., Li, Y., Takahashi, H. et al. (2015) Clonal analyses and gene profiling identify genetic biomarkers of the thermogenic potential of human brown and white preadipocytes. Nat Med, 21, 760–768.

78. Langston, S.P., Grossman, S., England, D., Afroze, R., Bence, N., Bowman, D., Bump, N., Chau, R., Chuang, B.-C., Claiborne, C. et al. (2021) Discovery of TAK-981, a First-in-Class Inhibitor of SUMO-Activating Enzyme for the Treatment of Cancer. Journal of Medicinal Chemistry, 64, 2501–2520.

79. Dudek, A.Z., Juric, D., Dowlati, A., Seymour, E.K., Ahnert, J.R., Wang, B.X., Huszar, D., Berger, A.J., Friedlander, S., Gomez-Pinillos, A. et al. (2021) Phase 1/2 study of the novel SUMOylation inhibitor TAK-981 in adult patients (pts) with advanced or metastatic solid tumors or relapsed/refractory (RR) hematologic malignancies. Journal of Clinical Oncology, 39.

80. Ohno, H., Shinoda, K., Spiegelman, B.M. and Kajimura, S. (2012) PPARgamma agonists induce a white-to-brown fat conversion through stabilization of PRDM16 protein. Cell Metab, 15, 395–404.

81. Bjune, J.-I., Laber, S., Lawrence-Archer, L., Nothnagel, P.M.C., Yamada, S., Zhao, X., Panahandeh Strømland, P., Al-Sharabi, N., Mustafa, K., Njølstad, P.R. et al. (2025) IRX3 controls a SUMOylation-dependent differentiation switch in adipocyte precursor cells. Nature Communications, 16, 7248.

82. Baig, M.S., Dou, Y., Bergey, B.G., Bahar, R., Burgener, J.M., Moallem, M., McNeil, J.B., Akhter, A., Burke, G.L., Sri Theivakadadcham, V.S. et al. (2021) Dynamic sumoylation of promoter-bound general transcription factors facilitates transcription by RNA polymerase II. PLoS Genet, 17, e1009828.

83. Launonen, K.-M., Varis, V., Aaltonen, N., Niskanen, E.A., Varjosalo, M., Paakinaho, V. and Palvimo, J.J. (2024) Central role of SUMOylation in the regulation of chromatin interactions and transcriptional outputs of the androgen receptor in prostate cancer cells. Nucleic Acids Research.

84. Chymkowitch, P., Nguea, A.P., Aanes, H., Koehler, C.J., Thiede, B., Lorenz, S., Meza-Zepeda, L.A., Klungland, A. and Enserink, J.M. (2015) Sumoylation of Rap1 mediates the recruitment of TFIID to promote transcription of ribosomal protein genes. Genome Res, 25, 897–906.

85. Chymkowitch, P., Nguea, P.A., Aanes, H., Robertson, J., Klungland, A. and Enserink, J.M. (2017) TORC1-dependent sumoylation of Rpc82 promotes RNA polymerase III assembly and activity. Proc Natl Acad Sci U S A, 114, 1039–1044.

86. Nguea, P.A., Robertson, J., Herrera, M.C., Chymkowitch, P. and Enserink, J.M. (2019) Desumoylation of RNA polymerase III lies at the core of the Sumo stress response in yeast. J Biol Chem, 294, 18784–18795.

87. Supek, F., Bošnjak, M., Škunca, N. and Šmuc, T. (2011) REVIGO Summarizes and Visualizes Long Lists of Gene Ontology Terms. PloS one, 6, e21800.

88. Guilherme, A., Yenilmez, B., Bedard, A.H., Henriques, F., Liu, D., Lee, A., Goldstein, L., Kelly, M., Nicoloro, S.M., Chen, M. et al. (2020) Control of Adipocyte Thermogenesis and Lipogenesis through β3-Adrenergic and Thyroid Hormone Signal Integration. Cell reports, 31, 107598.

89. Ellis, J.M., Li, L.O., Wu, P.-C., Koves, T.R., Ilkayeva, O., Stevens, R.D., Watkins, S.M., Muoio, D.M. and Coleman, R.A. (2010) Adipose Acyl-CoA Synthetase-1 Directs Fatty Acids toward &#x3b2;-Oxidation and Is Required for Cold Thermogenesis. Cell Metabolism, 12, 53–64.

90. Guilherme, A., Yenilmez, B., Bedard, A.H., Henriques, F., Liu, D., Lee, A., Goldstein, L., Kelly, M., Nicoloro, S.M., Chen, M. et al. (2020) Control of Adipocyte Thermogenesis and Lipogenesis through &#x3b2;3-Adrenergic and Thyroid Hormone Signal Integration. Cell reports, 31.

91. Goffeney, A., Hendriks, I.A., Morel, V., Loe-Mie, Y., Charon, F., Nielsen, M.L., Cossec, J.C., Noordermeer, D., Seeler, J.S. and Dejean, A. (2025) SUMO operates from a unique long tandem repeat to keep innate immunity in check. Nucleic Acids Res, 53.

92. Inagaki, T., Sakai, J. and Kajimura, S. (2016) Transcriptional and epigenetic control of brown and beige adipose cell fate and function. Nat Rev Mol Cell Biol, 17, 480–495.

93. Cao, W., Daniel, K.W., Robidoux, J., Puigserver, P., Medvedev, A.V., Bai, X., Floering, L.M., Spiegelman, B.M. and Collins, S. (2004) p38 mitogen-activated protein kinase is the central regulator of cyclic AMP-dependent transcription of the brown fat uncoupling protein 1 gene. Mol Cell Biol, 24, 3057–3067.

94. Wu, J., Bostrom, P., Sparks, L.M., Ye, L., Choi, J.H., Giang, A.H., Khandekar, M., Virtanen, K.A., Nuutila, P., Schaart, G. et al. (2012) Beige adipocytes are a distinct type of thermogenic fat cell in mouse and human. Cell, 150, 366–376.

95. Shinoda, K., Ohyama, K., Hasegawa, Y., Chang, H.-Y., Ogura, M., Sato, A., Hong, H., Hosono, T., Sharp, Louis Z., Scheel, David W. et al. (2015) Phosphoproteomics Identifies CK2 as a Negative Regulator of Beige Adipocyte Thermogenesis and Energy Expenditure. Cell Metabolism, 22, 997–1008.

96. Theurillat, I., Hendriks, I.A., Cossec, J.C., Andrieux, A., Nielsen, M.L. and Dejean, A. (2020) Extensive SUMO Modification of Repressive Chromatin Factors Distinguishes Pluripotent from Somatic Cells. Cell reports, 32, 108146.

97. Siersbæk, R., Nielsen, R. and Mandrup, S. (2010) PPARγ in adipocyte differentiation and metabolism – Novel insights from genome-wide studies. FEBS Letters, 584, 3242–3249.

98. Hendriks, I.A., Lyon, D., Su, D., Skotte, N.H., Daniel, J.A., Jensen, L.J. and Nielsen, M.L. (2018) Site-specific characterization of endogenous SUMOylation across species and organs. Nat Commun, 9, 2456.

99. Katafuchi, T., Holland, W.L., Kollipara, R.K., Kittler, R., Mangelsdorf, D.J. and Kliewer, S.A. (2018) PPARgamma-K107 SUMOylation regulates insulin sensitivity but not adiposity in mice. Proc Natl Acad Sci U S A, 115, 12102–12111.

100. Shao, M., Wang, Q.A., Song, A., Vishvanath, L., Busbuso, N.C., Scherer, P.E. and Gupta, R.K. (2019) Cellular Origins of Beige Fat Cells Revisited. Diabetes, 68, 1874–1885.

101. Roh, H.C., Tsai, L.T.Y., Shao, M., Tenen, D., Shen, Y., Kumari, M., Lyubetskaya, A., Jacobs, C., Dawes, B., Gupta, R.K. et al. (2018) Warming Induces Significant Reprogramming of Beige, but Not Brown, Adipocyte Cellular Identity. Cell Metab, 27, 1121–1137 e1125.

102. Decque, A., Joffre, O., Magalhaes, J.G., Cossec, J.C., Blecher-Gonen, R., Lapaquette, P., Silvin, A., Manel, N., Joubert, P.E., Seeler, J.S. et al. (2016) Sumoylation coordinates the repression of inflammatory and anti-viral gene-expression programs during innate sensing. Nat Immunol, 17, 140–149.

103. Siersbaek, R. and Mandrup, S. (2011) Transcriptional networks controlling adipocyte differentiation. Cold Spring Harb Symp Quant Biol, 76, 247–255.

104. Paakinaho, V., Lempiainen, J.K., Sigismondo, G., Niskanen, E.A., Malinen, M., Jaaskelainen, T., Varjosalo, M., Krijgsveld, J. and Palvimo, J.J. (2021) SUMOylation regulates the protein network and chromatin accessibility at glucocorticoid receptor-binding sites. Nucleic Acids Res, 49, 1951–1971.

105. González-Vinceiro, L., Espejo-Serrano, C., Soler-Oliva, M.E., Soto-Hidalgo, E., Mateos-Martín, M.L., Rico, D., González-Aguilera, C. and González-Prieto, R. (2025) PLAMseq enables the proteo-genomic characterization of chromatin-associated proteins and protein interactions in a single workflow. Science Advances, 11, eady4151.

106. Chen, Q., Huang, L., Pan, D., Zhu, L.J. and Wang, Y.-X. (2018) Cbx4 Sumoylates Prdm16 to Regulate Adipose Tissue Thermogenesis. Cell reports, 22, 2860–2872.

107. Zhao, J., Qian, H., An, Y., Chu, L., Tan, D., Qin, C., Sun, Q., Wang, Y. and Qi, W. (2024) PPAR&#x3b3; and C/EBP&#x3b1; enable adipocyte differentiation upon inhibition of histone methyltransferase PRC2 in malignant tumors. Journal of Biological Chemistry, 300.

108. Parreno, V., Loubiere, V., Schuettengruber, B., Fritsch, L., Rawal, C.C., Erokhin, M., Győrffy, B., Normanno, D., Di Stefano, M., Moreaux, J., et al. (2024) Transient loss of Polycomb components induces an epigenetic cancer fate. Nature.

109. Li, F., Wu, R., Cui, X., Zha, L., Yu, L., Shi, H. and Xue, B. (2016) Histone Deacetylase 1 (HDAC1) Negatively Regulates Thermogenic Program in Brown Adipocytes via Coordinated Regulation of Histone H3 Lysine 27 (H3K27) Deacetylation and Methylation*. Journal of Biological Chemistry, 291, 4523–4536.

110. Leonen, C.J.A., Shimada, M., Weller, C.E., Nakadai, T., Hsu, P.L., Tyson, E.L., Mishra, A., Shelton, P.M., Sadilek, M., Hawkins, R.D. et al. (2021) Sumoylation of the human histone H4 tail inhibits p300-mediated transcription by RNA polymerase II in cellular extracts. Elife, 10.

111. Yazıcı, E. and McIntyre, J. (2025) The complex network of p300/CBP regulation: Interactions, posttranslational modifications, and therapeutic implications. Journal of Biological Chemistry, 301, 110715.

112. Nyman, E., Bartesaghi, S., Melin Rydfalk, R., Eng, S., Pollard, C., Gennemark, P., Peng, X.-R. and Cedersund, G. (2017) Systems biology reveals uncoupling beyond UCP1 in human white fat-derived beige adipocytes. npj Systems Biology and Applications, 3, 29.

113. Aydemir, D., Sarayloo, E. and Nuray, U.N. (2020) Rosiglitazone-induced changes in the oxidative stress metabolism and fatty acid composition in relation with trace element status in the primary adipocytes. J Med Biochem, 39, 267–275.

114. Wang, Q., Xu, C., Fan, Q., Yuan, H., Zhang, X., Chen, B., Cai, R., Zhang, Y., Lin, M. and Xu, M. (2021) Positive feedback between ROS and cis-axis of PIASxα/p38α-SUMOylation/MK2 facilitates gastric cancer metastasis. Cell Death Dis, 12, 986.

115. Hendriks, I.A. and Vertegaal, A.C. (2016) A comprehensive compilation of SUMO proteomics. Nat Rev Mol Cell Biol, 17, 581–595.

116. Sri Theivakadadcham, V.S., Bergey, B.G. and Rosonina, E. (2019) Sumoylation of DNA-bound transcription factor Sko1 prevents its association with nontarget promoters. PLoS Genet, 15, e1007991.

117. Valima, E., Varis, V., Bureiko, K., Lempiäinen, J.K., Schroderus, A.M., Oksa, L., Lohi, O., Kinnunen, T., Varjosalo, M., Niskanen, E.A. et al. (2025) SUMOylation inhibition potentiates the glucocorticoid receptor to program growth arrest of acute lymphoblastic leukemia cells. Oncogene.

118. Hartig, S.M., Bader, D.A., Abadie, K.V., Motamed, M., Hamilton, M.P., Long, W., York, B., Mueller, M., Wagner, M., Trauner, M. et al. (2015) Ubc9 Impairs Activation of the Brown Fat Energy Metabolism Program in Human White Adipocytes. Mol Endocrinol, 29, 1320–1333.

119. Lee, J.S., Chae, S., Nan, J., Koo, Y.D., Lee, S.A., Park, Y.J., Hwang, D., Han, W., Lee, D.S., Kim, Y.B. et al. (2022) SENP2 suppresses browning of white adipose tissues by de-conjugating SUMO from C/EBPbeta. Cell reports, 38, 110408.

