## Supplementary files for "Transient SUMOylation Inhibition In Human Pre-adipocytes Stably Imprints a Transcriptional Beiging Fate"

**Supplementary material**

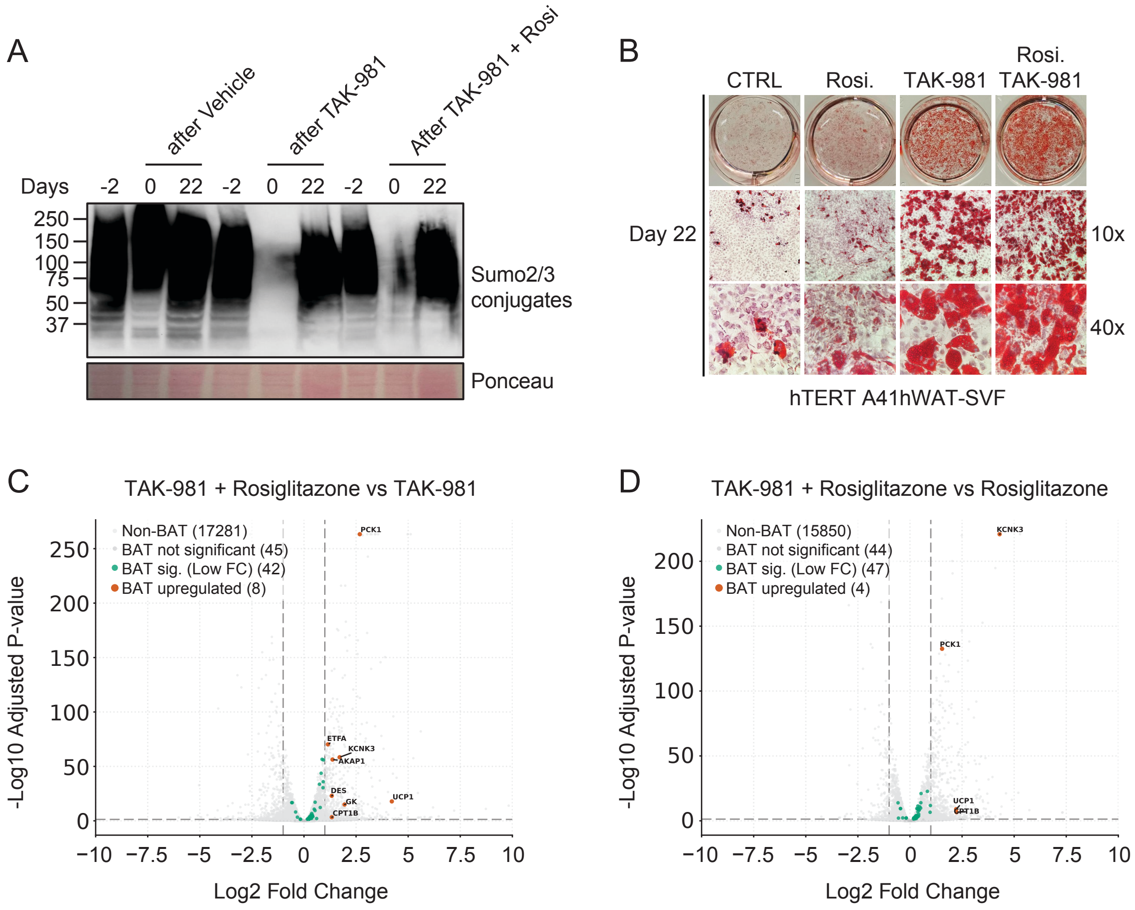

**Suppl. Fig. 1:**

(A) Assessment of the effect of Rosiglitazone on SUMOylation during adipogenesis.

(B) Oil-Red-O staining assessment of fat accumulation 22 days after adipogenic induction in hTERT A41hWAT-SVF pre-adipocytes upon TAK-981 and rosiglitazone treatments.

(C-D) Volcano plots displaying differentially expressed genes (DEGs; $log_2FC > 1$ or $log_2FC < -1$; padj < 0.05) resulting from co-treatment with TAK-981 + Rosiglitazone compared to TAK-981 *(C)* or Rosiglitazone *(D)*. Data were collected in mature adipocytes 21 days post adipogenic induction. Significant BATLAS DEGs are highlighted.

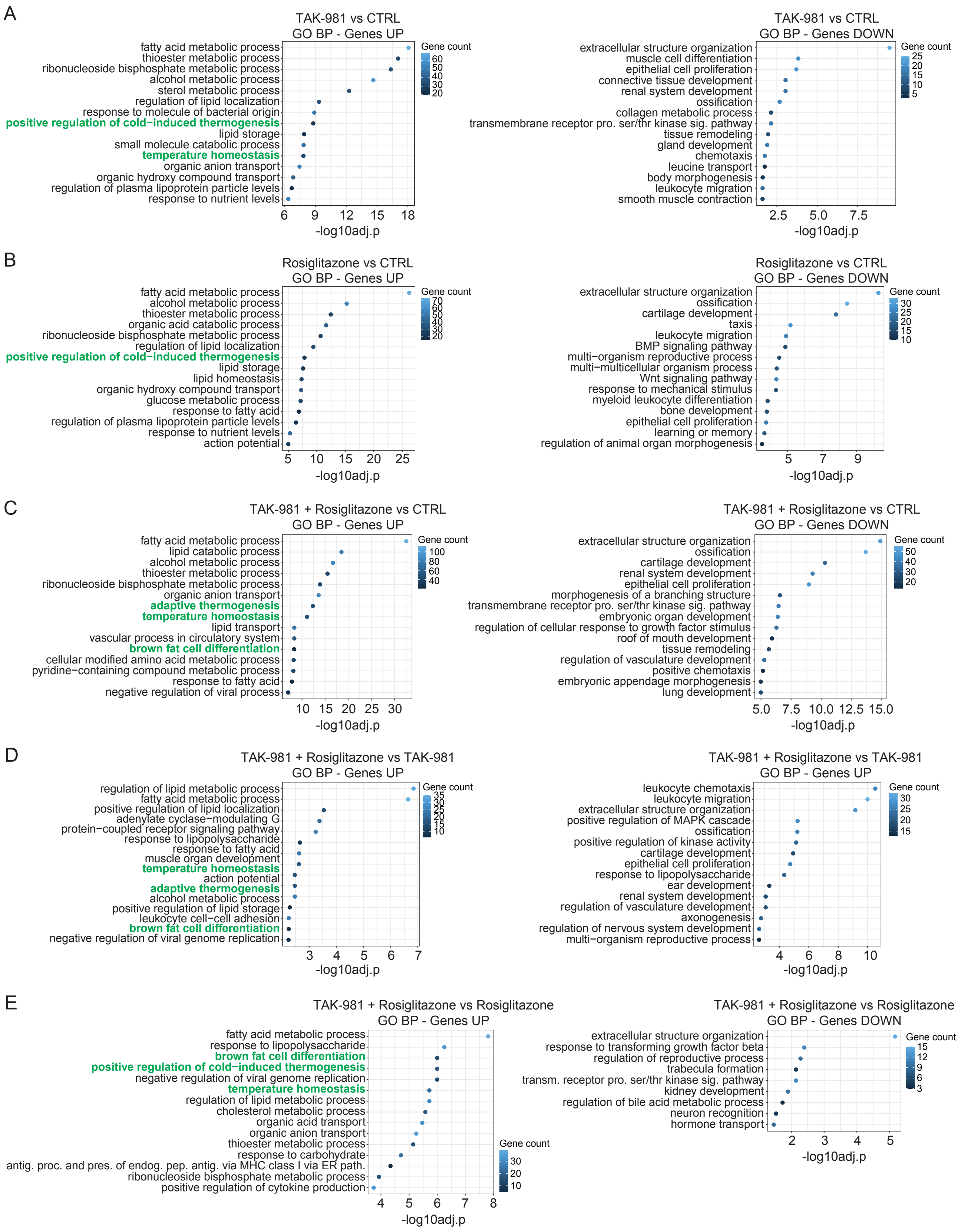

**Suppl. Fig. 2**

(A) Gene Ontology (GO) analysis of Biological Processes associated with the DEGs in Fig. 1C.

(B) Gene Ontology (GO) analysis of Biological Processes associated with the DEGs in Fig. 1D.

(C) Gene Ontology (GO) analysis of Biological Processes associated with the DEGs in Fig. 1E.

(D) Gene Ontology (GO) analysis of Biological Processes associated with the DEGs in Suppl. Fig. 1C.

(E) Gene Ontology (GO) analysis of Biological Processes associated with the DEGs in Suppl. Fig. 1D.

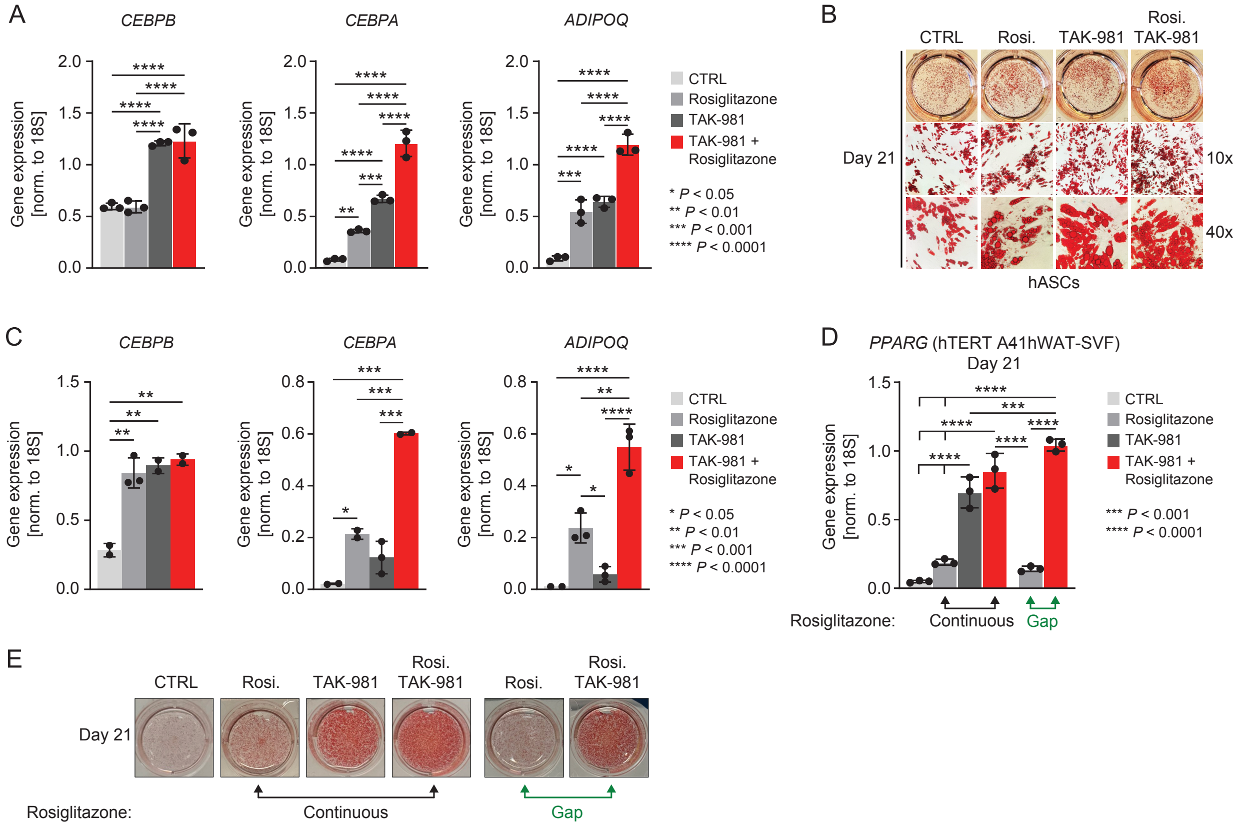

**Suppl. Fig. 3**

(A) RT-qPCR analysis of adipogenic marker 22 days post adipogenic induction in hTERT A41hWAT-SVF pre-adipocytes following TAK-981 and rosiglitazone treatments (see Fig. 1A). Data were normalized to the expression of *18S* and represent the mean ± standard deviation from at least three independent experiments.

(B) Oil-Red-O staining assessment of fat accumulation 22 days after adipogenic induction in human adipose stem cells (hASCs) upon TAK-981 and rosiglitazone treatments.

(C) RT-qPCR analysis of adipogenic marker 22 days post adipogenic induction in hASCs following TAK-981 and rosiglitazone treatments. Data were normalized to the expression of *18S* and represent the mean ± standard deviation from at least three independent experiments.

(D) RT-qPCR analysis of *PPARG* mRNA expression in hTERT A4hWAT-SVF cells under the conditions described in Fig. 2G. Data were normalized to the expression of *18S* and represent the mean ± standard deviation from at least three independent experiments.

(E) Oil-Red-O staining assessment of fat accumulation 22 days after adipogenic induction in hTERT A4hWAT-SVF cells upon TAK-981 and rosiglitazone treatments under condition described in Fig. 2G.

Statistical analyses: ONE-WAY-ANOVA with Tukey's multiple comparison test.

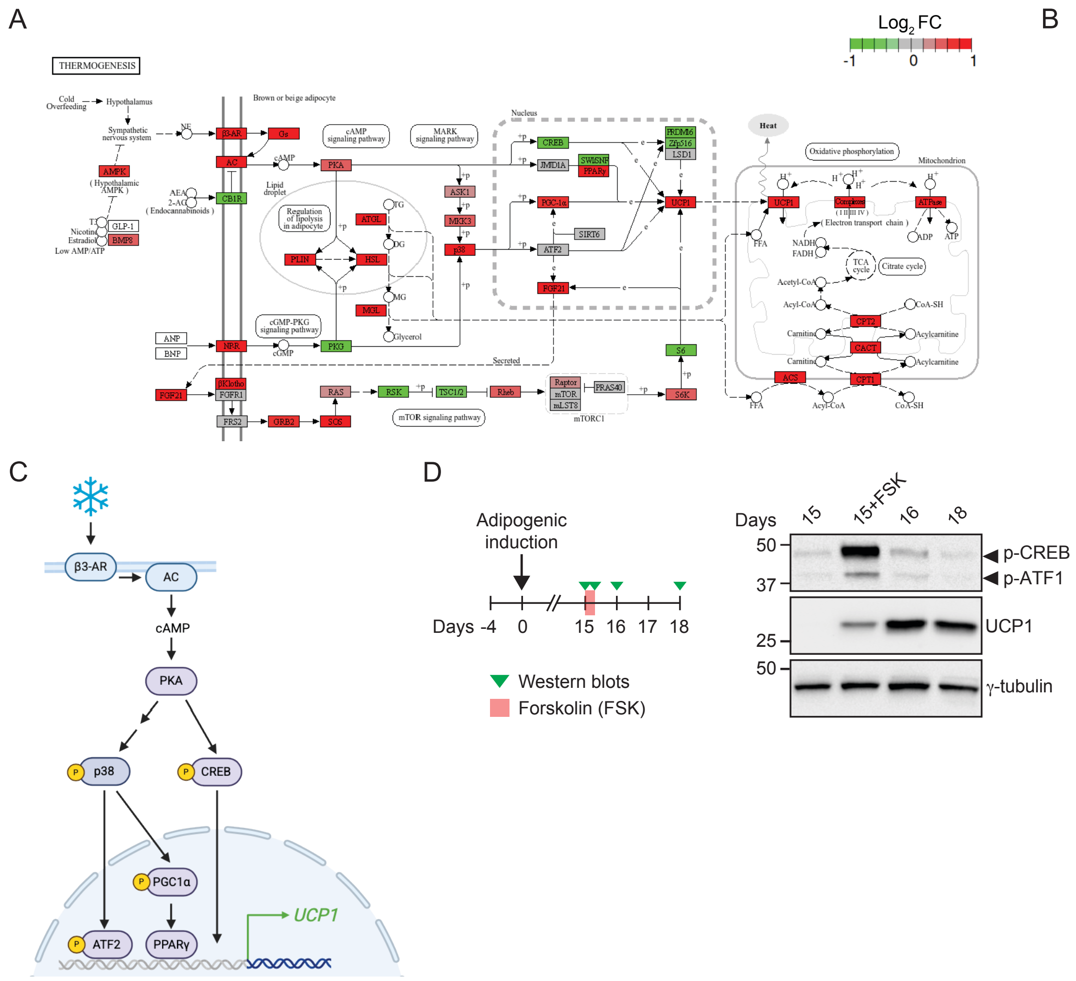

**Suppl. Fig. 4**

(A) The “Thermogenesis” KEGG pathway was visualized using differentially expressed genes between TAK-981 + Rosiglitazone and control cells. Pathway visualization was performed using the pathview (v1.44.0) R package.

(B) Hypothesis: SUMOylation in pre-adipocytes blunts the cAMP-PKA-p38 signaling pathway in mature adipocytes, limiting adaptive thermogenesis.

(C) Simplified view of adaptive thermogenesis displaying the phosphor-substrates analyzed in Fig. 3. Diagram produced using Biorender®.

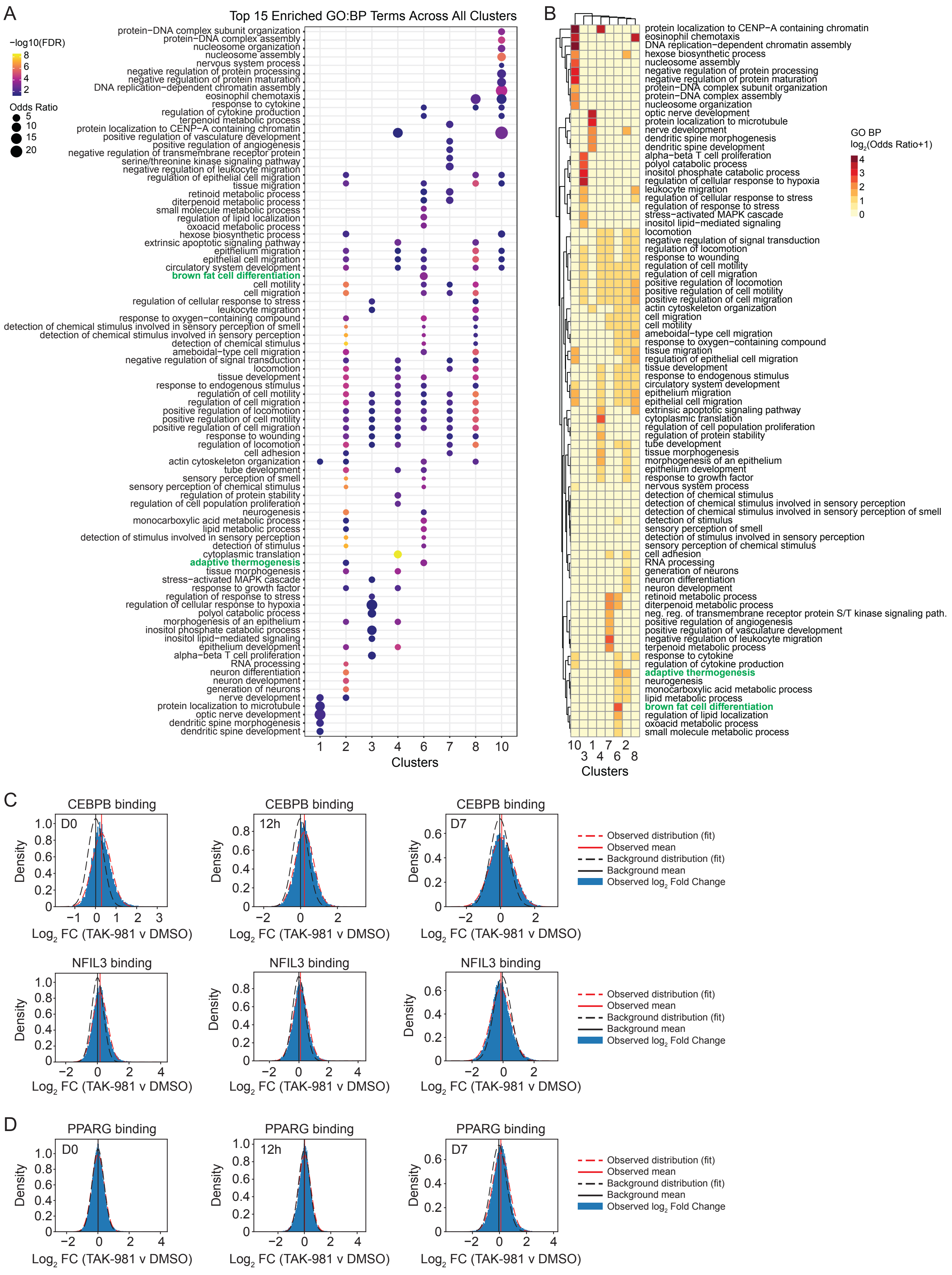
A
Top 15 Enriched GO:BP Terms Across All Clusters
−log10(FDR)
Odds Ratio
Clusters
B
GO BP
log2(Odds Ratio+1)
Clusters
C
CEBPB binding
NFIL3 binding
Density
Log2 FC (TAK-981 v DMSO)
Observed distribution (fit)
Observed mean
Background distribution (fit)
Background mean
Observed log2 Fold Change
D
PPARG binding

**Suppl. Fig. 5:**

(A) Dot plot summarizing the enrichment of top Gene Ontology Biological Process (GO:BP) terms across all clusters. The terms shown are a union set representing the top 15 most significant (by FDR, then Odds Ratio) terms from each individual cluster. The size of each point corresponds to the enrichment magnitude (Odds Ratio), while the color intensity reflects statistical significance (-log10 FDR).
(B) Heatmap displaying the enrichment profiles for the set of GO:BP terms from *(A)*. Cell color represents the enrichment magnitude, calculated as the log2-transformed Odds Ratio (log2(Odds Ratio + 1)). Both terms (rows) and the clusters (columns) are hierarchically clustered to group similar profiles.
(C-D) Density plots showing the timely differential mobilization (TAK-981 vs DMSO) of selected TFs CEBPB, NILF3 *(C)* and PPARG *(D)* at day 0, 12 hours and day 7 (see experimental layout in Fig. 5A).

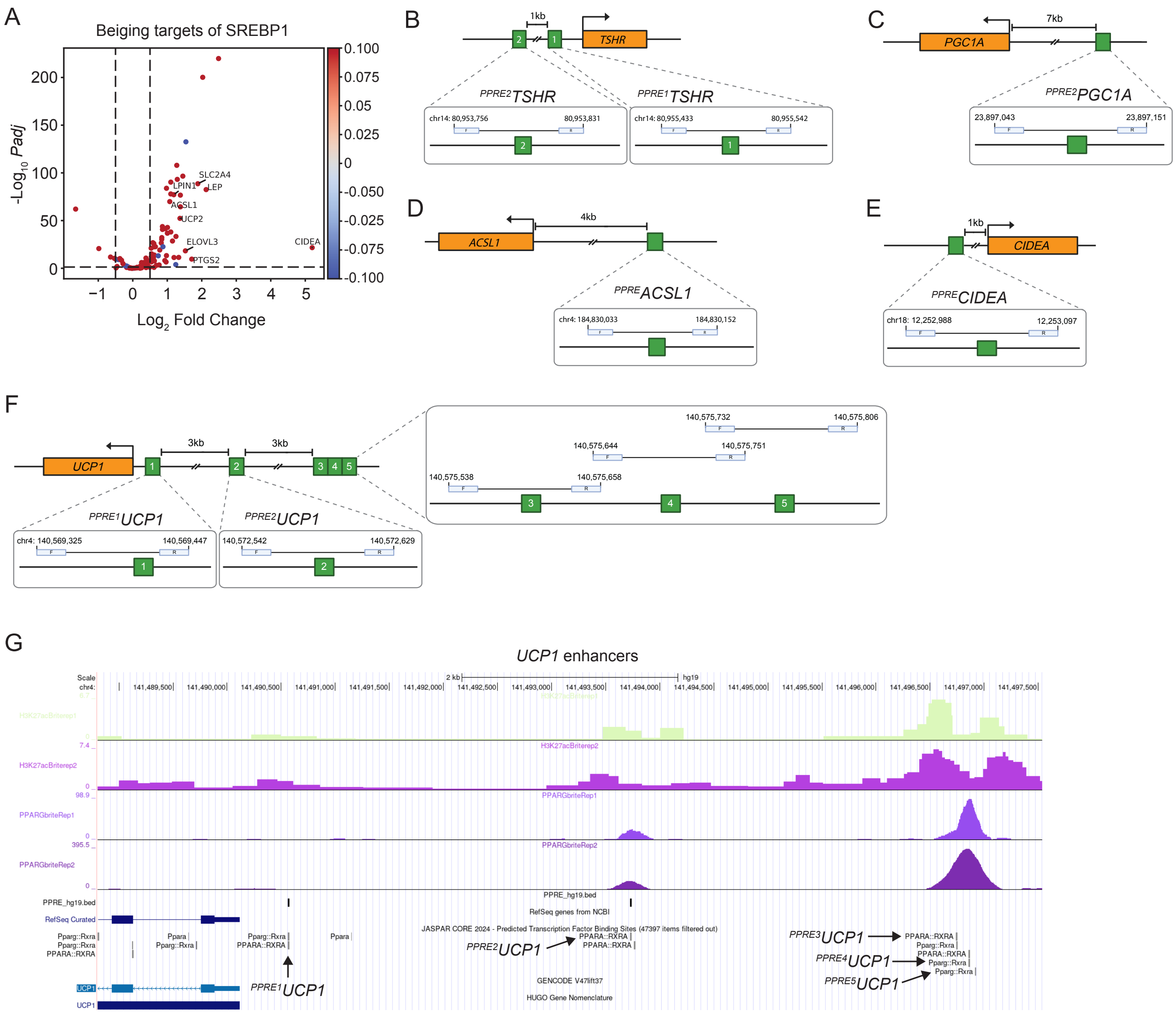

**Suppl. Fig. 6:**

(A) Volcano plot showing SREBP1-target genes activated in response to transient SUMOylation inhibition and rosiglitazone. DEGs were identified by comparing TAK-981 + Rosiglitazone-treated cells to Rosiglitazone-treated cells.

(B-F) Schematics of ChIP-qPCR regions and primer pairs used in the experiments in Fig. 6D-F.

(G) Genome track generated using the UCSC genome browser showing the presence of PPARG and H3K27ac at *UCP1* enhancers (studied in Fig. 6D,F).

**Supplementary Tables 2, 3, 4, 5 and 6** have been uploaded separately.

**Supplementary Table 1: Oligonucleotides used in this study.**

| Oligonucleotide | Sequence (5′-3′) | Use |
|---|---|---|
| **18S_F** | GCAGAATCCACGCGCCAGTACAAG | RT-qPCR |
| **18S_R** | GCTTGTTGTCCAGACCATTGGC | |
| **CEBPA_F** | GACTTGGTGCGTCTAAGATGAG | |
| **CEBPA_R** | GGCATTGGAGCGGTGAGTT | |
| **CEBPB_F** | AGAAGACCGTGGACAAGCACAG | |
| **CEBPB_R** | CTCCAGGACCTTGTGCTGCGT | |
| **PPARG2_F** | ACTCTGGGAGATTCTCCTATT | |
| **PPARG2_R** | CTCCATAGTGAAATCCAGAAG | |
| **ADIPOQ_F** | CAGGCCGTGATGGCAGAGATG | |
| **ADIPOQ_R** | GGTTTCACCGATGTCTCCCTTAG | |
| **UCP1_F** | ACCGCAGGGAAAGAAACAGC | |
| **UCP1_R** | TCAGATTGGGAGTAGTCCCT | |
| **CTRL_F** | GCCCTCCTGGGATTAAA | ChIP-qPCR |
| **CTRL_R** | CATTCCATTCCCACGAAC | |
| **hUCP1_PPRE1_F** | GCTCTCCCAGAAGGAAG | |
| **hUCP1_PPRE1_R** | GAGGAAGGAGGATGAGAAG | |
| **hUCP1_PPRE2_F** | CTGGCTTCACCACTTCT | |
| **hUCP1_PPRE2_R** | GGGATACCACCCTCTCC | |
| **hUCP1_PPRE3_F** | CTGGTATGCCGTGACTAT | |
| **hUCP1_PPRE3_R** | GTCCGGGAGATTAGAAAGA | |
| **hUCP1_PPRE4_F** | TCTAATCTCCCGGACAAAT | |
| **hUCP1_PPRE4_R** | TGTCTGCCTCACATTCA | |
| **hUCP1_PPRE5_F** | GAGTGAATGTGAGGCAGA | |
| **hUCP1_PPRE5_R** | CATCACAAAGAATGCTTCCA | |
| **hPGC1A_PPRE1_F** | TGACTTGTGTTTCTGTTATGG | |
| **hPGC1A_PPRE1_R** | CTACCTCTCCGACATGTTT | |
| **hPGC1A_PPRE2_F** | GGCTTCTCAAATATCACAAGAG | |
| **hPGC1A_PPRE2_R** | GAGAACATGAAATCAAATTGGC | |
| **hCIDEA_PPRE_F** | CAGCCCTGTTCTTTCTATG | |
| **hCIDEA_PPRE_R** | CCCAATGTGATGGATGAG | |
| **hTSHR_PPRE1_F** | CCAGCCAAGGAAAGTGAA | |
| **hTSHR_PPRE1_R** | TCACCACTGTGGAGGAG | |
| **hTSHR_PPRE2_F** | TTCTCTGGTGTTCATTTGTAG | |
| **hTSHR_PPRE2_R** | AAACCATAGGGACAGGAAA | |
| **hACSL1_PPRE_F** | GCAGTGAACCGTGATTG | |
| **hACSL1_PPRE_R** | TCAGCTTGACCCAGAAA | |
| **Ad1_noMX** | AATGATACGGCGACCACCGAGATCTACACTCGTCGGCAGCGTCAGATGTG | ATAC-seq |
| **Ad2.1_TAAGGCGA** | CAAGCAGAAGACGGCATACGAGATTCGCCTTAGTCTCGTGGGCTCGGAGATGT | |
| **Ad2.2_CGTACTAG** | CAAGCAGAAGACGGCATACGAGATCTAGTACGGTCTCGTGGGCTCGGAGATGT | |
| **Ad2.3_AGGCAGAA** | CAAGCAGAAGACGGCATACGAGATTTCTGCCTGTCTCGTGGGCTCGGAGATGT | |
| **Ad2.4_TCCTGAGC** | CAAGCAGAAGACGGCATACGAGATGCTCAGGAGTCTCGTGGGCTCGGAGATGT | |

| | |
|---|---|
| **Ad2.5_GGACTCCT** | CAAGCAGAAGACGGCATACGAGATAGGAGTCCGTCTCGTGGGCTCGGAGATGT |
| **Ad2.6_TAGGCATG** | CAAGCAGAAGACGGCATACGAGATCATGCCTAGTCTCGTGGGCTCGGAGATGT |
| **Ad2.7_CTCTCTAC** | CAAGCAGAAGACGGCATACGAGATGTAGAGAGGTCTCGTGGGCTCGGAGATGT |
| **Ad2.8_CAGAGAGG** | CAAGCAGAAGACGGCATACGAGATCCTCTCTGGTCTCGTGGGCTCGGAGATGT |
| **Ad2.9_GCTACGCT** | CAAGCAGAAGACGGCATACGAGATAGCGTAGCGTCTCGTGGGCTCGGAGATGT |
| **Ad2.10_CGAGGCTG** | CAAGCAGAAGACGGCATACGAGATCAGCCTCGGTCTCGTGGGCTCGGAGATGT |
| **Ad2.11_AAGAGGCA** | CAAGCAGAAGACGGCATACGAGATTGCCTCTTGTCTCGTGGGCTCGGAGATGT |
| **Ad2.12_GTAGAGGA** | CAAGCAGAAGACGGCATACGAGATTCCTCTACGTCTCGTGGGCTCGGAGATGT |
| **Ad2.13_GTCGTGAT** | CAAGCAGAAGACGGCATACGAGATATCACGACGTCTCGTGGGCTCGGAGATGT |
| **Ad2.14_ACCACTGT** | CAAGCAGAAGACGGCATACGAGATACAGTGGTGTCTCGTGGGCTCGGAGATGT |
| **Ad2.15_TGGATCTG** | CAAGCAGAAGACGGCATACGAGATCAGATCCAGTCTCGTGGGCTCGGAGATGT |
